# Cholesterol-Responsive Interaction of NFE2L1-INSIG1 Controls VLDL Secretion and MASH Pathogenesis

**DOI:** 10.1101/2025.06.09.657856

**Authors:** Shijun Deng, Jessica Freed, Grace Y. Lee, Gizel Askin, Zhe Cao, Özgür Cakici, Bo Yuan, Sheng Hui, Karen E. Inouye, Isabel Graupera, Gökhan S. Hotamisligil

## Abstract

Cholesterol overload drives metabolic dysfunction-associated steatohepatitis (MASH). While the liver maintains homeostasis by exporting cholesterol via very low-density lipoprotein (VLDL) secretion, this process is paradoxically suppressed under cholesterol excess by insulin-induced gene 1 (INSIG1), which inhibits sterol regulatory element-binding protein 1 (SREBP1) activity. How VLDL secretion persists during cholesterol overload to prevent lipotoxicity remains unresolved. We identify a cholesterol-responsive interaction between nuclear factor erythroid 2 related factor-1 (NFE2L1) and INSIG1 that sustains cholesterol balance. Hepatic NFE2L1 deficiency elevates INSIG1 levels, suppressing SREBP1 activation and impairing VLDL secretion, leading to cholesterol accumulation and liver injury. Mechanistically, NFE2L1 binds to INSIG1 via its N-terminal homology box 2 (NHB2) domain, with cholesterol enhancing this interaction to drive INSIG1 degradation and SREBP1 activation. In NFE2L1-deficient mice, wild-type NFE2L1 restores SREBP1 activity and VLDL secretion, while the NHB2-deleted mutant (ΔNHB2) fails. Lipidomics reveal that NFE2L1 deficiency reduces serum triglyceride composition, restored exclusively by wild-type NFE2L1. In a murine MASH model, NFE2L1 overexpression activates SREBP1/2, enhances cholesterol secretion, and alleviates liver injury, inflammation and fibrosis, without elevating atherogenic lipoproteins due to compensatory LDL receptor upregulation. Our findings resolve the paradox of cholesterol-driven VLDL secretion and establish the NFE2L1-INSIG1 axis as a therapeutic target for metabolic diseases.

## INTRODUCTION

Metabolic dysfunction-associated steatohepatitis (MASH) affects over 5% of the global population, yet therapeutic strategies remain limited due to incomplete understanding of hepatic cholesterol homeostasis, a central driver of disease progression [1]. Cholesterol balance is critial for metabolic health, sustaining cellular functions, lipid metabolism, and physiological balance. Both low and high cholesterol levels disrupt cellular function and viability, necessitating adaptive mechanisms to coordinate responses to cholesterol fluctuations. The liver is integral to cholesterol synthesis, storage, and secretion through very-low-density lipoproteins (VLDL) [2, 3]. Dysregulated VLDL secretion contributes to MASH, hypercholesterolemia, and cardiovascular diseases by promoting hepatic lipid buildup and systemic cholesterol imbalance [4–6].

The sterol regulatory element-binding protein (SREBP) pathway orchestrates lipid biosynthesis and VLDL secretion, preventing hepatic lipid accumulation and maintaining metabolic homeostasis [7, 8]. SREBPs are a family of basic helix-loop-helix leucine zipper (bHLH-LZ) transcription factors that includes SREBP1 and SREBP2. SREBP1 promotes the expression of genes involved in fatty acids and triglyceride (TG) synthesis and increases microsomal triglyceride transfer protein (MTP) levels, which is essential for VLDL secretion [9]. By supplying lipid substrates needed for VLDL formation, SREBP1 also contributes to systemic lipid homeostasis [10]. SREBP1 activation is regulated by the interaction between insulin-induced gene 1 (INSIG1) and SREBP cleavage-activating protein (SCAP) in the endoplasmic reticulum (ER) [11]. Under low sterol conditions, INSIG1 dissociates from the SCAP/SREBP complex, enabling its transport to the Golgi. This releases the transcriptionally active N-terminal domain of SREBP1 [SREBP1(N)], which enters the nucleus to activate lipogenic genes. Conversely, high sterol levels stabilize INSIG1-SCAP binding, retaining SREBP1 in the ER and halting lipid synthesis [12, 13]. While this feedback loop prevents lipid overload, how SREBP1 remains functional to sustain VLDL secretion during cholesterol overload remains unresolved.

Nuclear factor erythroid-2 like 1 (NFE2L1/NRF1) has emerged as a critical regulator of hepatic cholesterol homeostasis, particularly in responding to increased intracellular cholesterol levels [14]. This Cap’N’Collar (CNC) transcription factor is also integrated into the ER membrane and influences proteasome expression as well as oxidative stress responses [15, 16]. Under certain stress conditions, NFE2L1 can be cleaved and relocate to the nucleus, suggesting that it may coordinate both transcription-dependent and non-transcriptional regulatory functions. In its ER-resident form, NFE2L1 directly senses free cholesterol (FC) to promote cholesterol export, and its deficiency leads to hepatic cholesterol accumulation under dietary high-cholesterol challenge [14, 17]. However, if and how these two critical pathways, NFE2L1 and SREBP, responding to high and low cholesterol levels, respectively, intersect with each other to control fluctuations in cholesterol levels and coordinate proper cellular responses remain an important but unresolved question. Clarifying this connection could yield a more comprehensive understanding of cholesterol homeostasis and its impact on metabolic health.

Here, we identify a direct interaction between NFE2L1 and INSIG1 proteins that regulates SREBP1 activity, and consequently, VLDL secretion from liver. This interaction balances cholesterol retention and VLDL assembly, uncovering a new mechanism by which hepatic lipid homeostasis is maintained across cholesterol levels. Furthermore, we show that NFE2L1 expression protects against MASH by promoting cholesterol secretion and redistribution into the HDL fraction and attenuates liver inflammation and fibrogenesis. Our findings unify two pivotal cholesterol-regulatory frameworks, positioning the NFE2L1-INSIG1 axis as a potential target for metabolic disorders.

## RESULTS

### Hepatic NFE2L1-deficiency reduces SREBP1 activity and impairs lipid secretion

To investigate whether NFE2L1 regulates SREBP1 activity, we transiently suppressed NFE2L1 in cultured hepa1-6 hepatocytes using siRNA. Depletion of NFE2L1 selectively reduced levels of the transcriptionally active form SREBP1(N) (65 kDa), while the precursor SREBP1(P) (130 kDa) remained unchanged (**Fig. 1A-B**). Notably, SREBP2 protein levels could not be assessed due to the lack of a validated antibody for murine SREBP2. These results demonstrate that NFE2L1 is required for SREBP1 activity *in vitro*.

**Figure 1.**
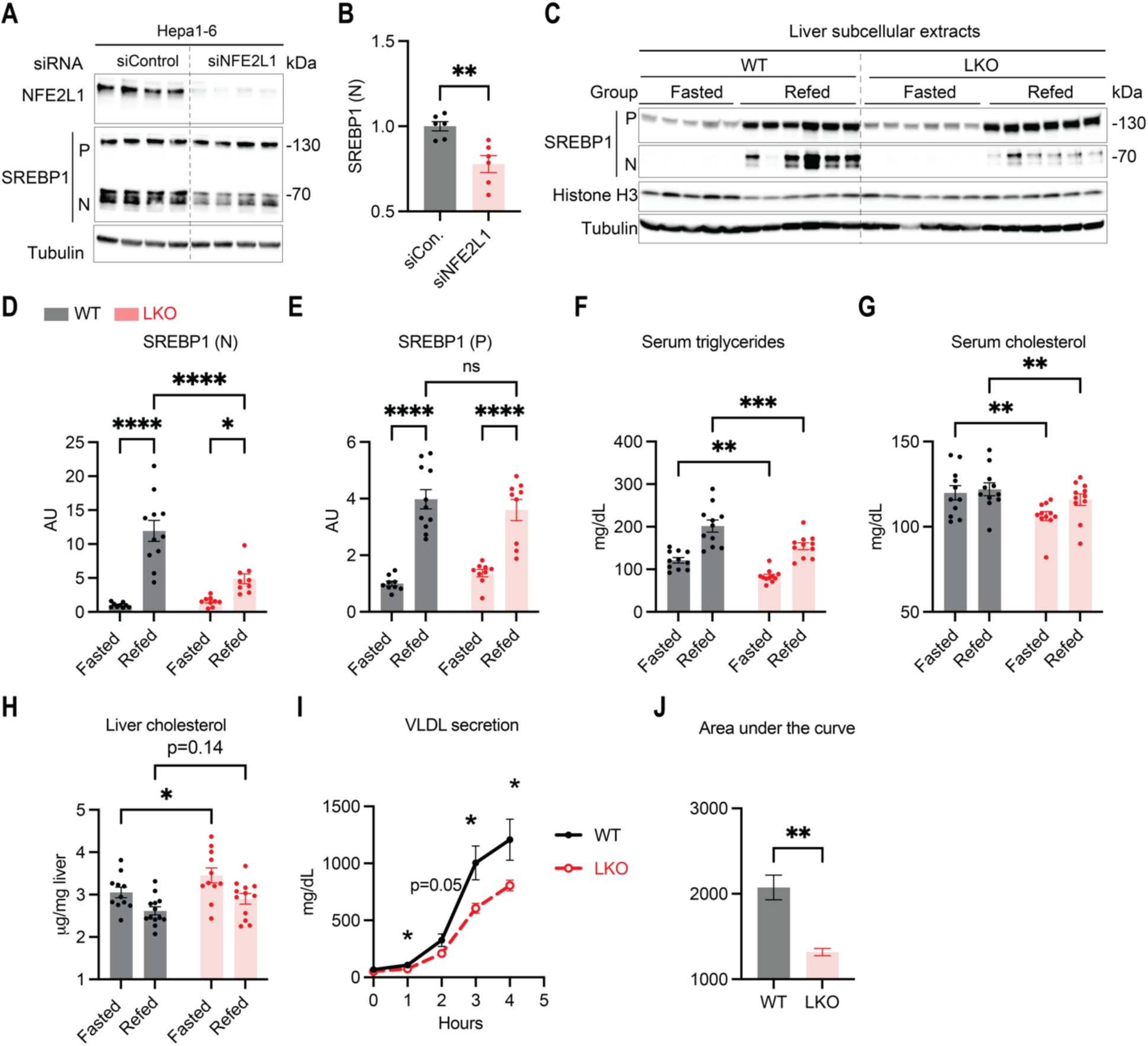
Hepatic NFE2L1-deficiency impairs SREBP1 activity and systemic lipid homeostasis. **(A-B)** Immunoblot analysis and quantification of SREBP1 in Hepa1-6 hepatocytes after transcient siRNA-mediated NFE2L1 knockdown (48 hours). **(C)** Subcellular fractionation of livers from 8-week-old liver-specific NFE2L1 knockout (LKO) and WT littermates under fasting or refed conditions (n=11/ group). Cytosolic and nuclear fraction of proteins were prepared from individual livers. Immunoblots show SREBP1(N) in the nuclear fractions and SREBP1(P) in the cytosolic fractions. **(D-E)** Quntification of SREBP1(N) (normalized to histone H3) and SREBP1(P) (normalized to tubulin) from (C). **(F-G)** Serum triaglyceride (TG) and total cholesterol in fasted/refed WT and LKO mice. **(H)** Liver total cholesterol in fasted/refed WT and LKO mice. **(I-J)** VLDL secretion: Plasma TG accumulation after tyloxapol injection in fasted WT and LKO mice (n=5-8/group). Data are means ± SEM. Statistics: *t*-test or two-way ANOVA (*p<0.05, **p<0.01, ***p<0.001,****p<0.0001).

We next tested whether hepatocyte-specific NFE2L1 knockout (LKO) mice exhibit defective SREBP1 processing *in vivo*. Since SREBP1 activation is dynamically regulated by nutritional status [18], wild-type (WT) and LKO mice were subjected to either overnight fasting (fasted) or fasting followed by a 6-hour refeeding period (refed). While LKO mice showed no differences in food intake, body weight, and serum levels of non-esterified fatty acids (NEFA) **(Fig. S1A-C)**, subcellular fractionation of liver tissue revealed striking defects in SREBP1 activation. In the WT mice, refeeding robustly increased nuclear SREBP1(N) levels (65 kDa) compared to fasting, consistent with feeding-driven SREBP1 cleavage. In contrast, LKO exhibited more than 50% reduction in nuclear SREBP1(N) under refed conditions (*p* < 0.0001 vs. refed WT, **Fig. 1C-D**), the precursor SREBP1(P) (130 kDa, ER membrane-bound) remained unchanged (**Fig. 1E**). This indicates that NFE2L1 deficiency impairs proteolytic activation and nuclear translocation of SREBP1. Consistent with the reduced SREBP1(N), mRNA levels of SREBP1/2 target genes, *Fasn*, *Acly*, *Ldlr*, *Hmgcr, Pcsk9* and *Fdps*, were markedly downregulated in LKO livers (**Fig. 2S**), despite unaltered SREBP1 transcription. These results establish that NFE2L1 is essential for feeding-stimulated SREBP1 processing and subsequent transcriptional activation of lipogenic program in the liver.

**Figure 2.**
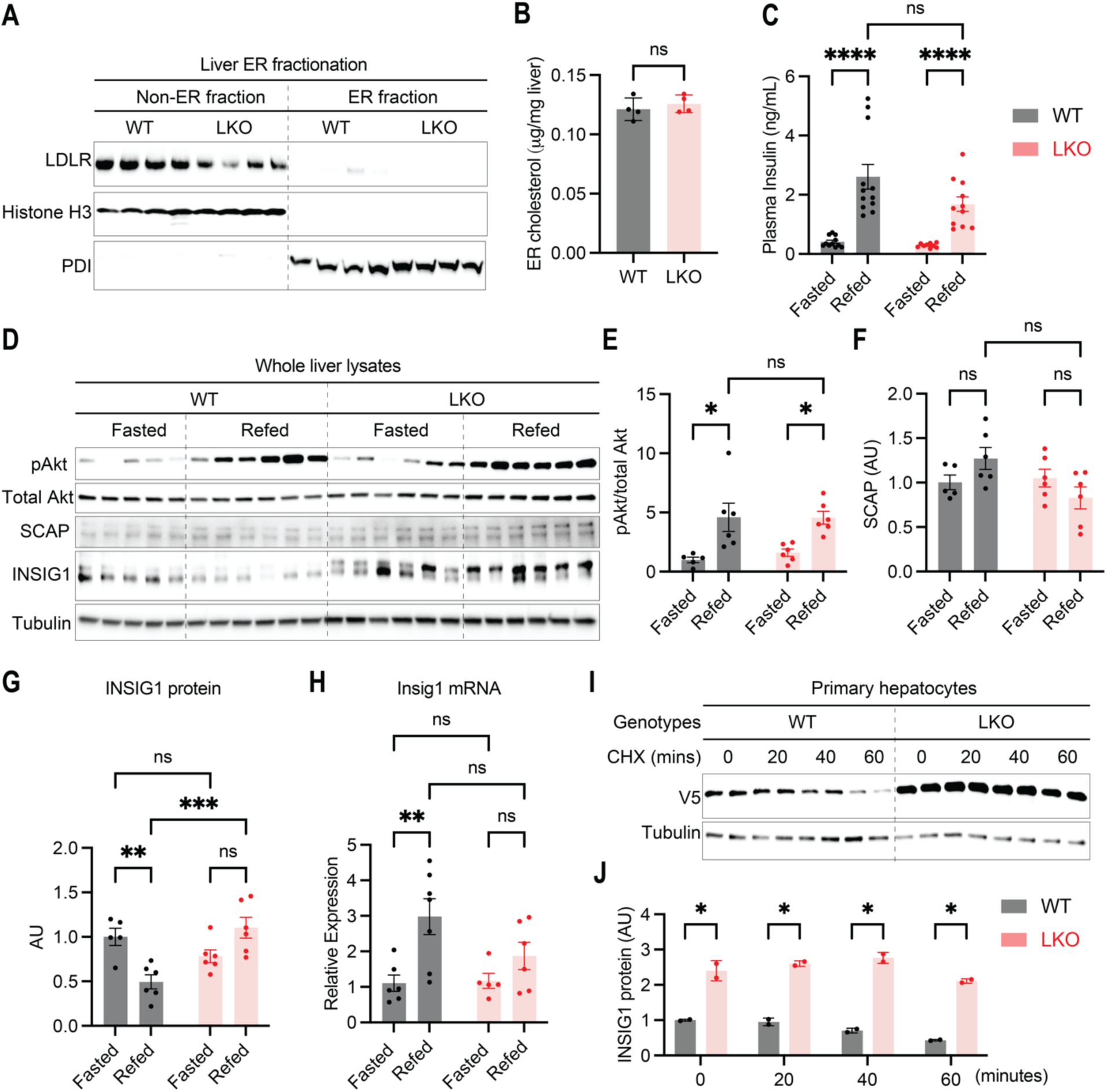
Hepatic NFE2L1 deficiency stabilizes INSIG1 protein. **(A)** ER-enriched fraction validation in livers of refed WT and LKO mice (n=4/group). Blots were probed for subcellular markers: LDLR (plasma membrane), Histone H3 (nuclear), and PDI (ER). **(B)** Cholesterol levels in the ER fractions isolated in (A). **(C)** Serum insulin levels in fasted/refed WT and LKO mice (n=11/group). **(D)** Immunoblot analysis of liver lysates from fasted/refed WT and LKO mice (n=5-6/group). **(E-G)** Quantification of pAkt/total Akt, SCAP, and INSIG1 normalized to Tubulin. **(H)** qRT-PCR analysis of *Insig1* mRNA in liver tissues from (D). **(I-J)** Cycloheximide (CHX) chase assay in primary hepatocytes isolated from WT and LKO mice injected with AAV-INSIG1-V5. Primary hepatocytes were treated with 10 µg/ml CHX for 0, 20, 40, or 60 minutes (n=2/conditions). Immunoblots are representative of 2 independent experiments. Data: Mean ± SEM. Statistcs: *t*-test or two-way ANOVA (*p<0.05, **p<0.01, ***p<0.001,****p<0.0001).

Given the central role of SREBP1 in lipid homeostasis, we next assessed systemic and hepatic lipids in the LKO mice. Liver-specific NFE2L1 deficiency reduced serum triglycerides (TG) and total cholesterol in both fasted and refed state (**Fig. 1F-G**), while liver TG remained unchanged (**Fig. S1D**). Strikingly, hepatic cholesterol content was significantly elevated in the fasted LKO mice (**Fig. 1H**), suggesting defective cholesterol export. After fasting most serum lipids are derived from hepatic VLDL secretion, hence, these results support that VLDL biogenesis and secretion maybe impaired in the livers of LKO mice. To directly test this, we blocked peripheral TG hydrolysis with tyloxapol (a lipoprotein lipase inhibitor) and quantified TG accumulation in the plasma overtime, which reflect the rate of VLDL secretion [19]. In WT mice, plasma TG rose linearly, reflecting robust VLDL secretion. In contrast, LKO exhibited a significant reduction in TG accumulation rate (**Fig. 1I-J**), confirming defective VLDL secretion. Together, these findings establish that NFE2L1 deficiency disrupts SREBP1-mediated VLDL assembly and secretion.

### NFE2L1 deficiency increases INSIG1 and suppress SREBP1 activation

To elucidate how hepatic NFE2L1 regulates SREBP1, we next evaluate factors controlling SREBP1 nuclear translocation, including cholesterol content in the ER, insulin signaling, and SCAP and INSIG1 protein stability. We isolated ER membranes from liver tissue and assessed ER cholesterol content. ER-enriched fraction revealed the presence of the ER-resident marker protein disulfide isomerase (PDI), while plasma membrane marker low density lipoprotein receptor (LDLR) and nuclear marker Histone H3 were undetectable, confirming the purity of the ER isolation **(Fig. 2A)**. ER-enriched fractions from refed mice showed comparable cholesterol levels between WT and LKO mice **(Fig. 2B)**. Likewise, serum insulin levels and hepatic insulin signaling (as indicated by pAkt levels), and SCAP protein levels were comparable between genotypes (**Fig. 2C-F**), ruling out these factors as primary causes of altered SREBP1 activity. Strikingly, fasting-refeeding regulation of INSIG1 protein was abolished in the LKO livers, with a 2-fold increase of endogenous INSIG1 protein in refed LKO livers **(Fig. 2G**). Since INSIG1 is a known SREBP1 target, we assessed whether this upregulation was transcriptionally driven by measuring *Insig1* mRNA. Despite blunted refeeding-induced *Insig1* mRNA upregulation in LKO livers, *Insig1* mRNA levels did not differ significantly between refed WT and LKO **(Fig. 2H).** This disconnection between mRNA and proteins indicates that elevated INSIG1 protein in LKO livers may arise from a post-translational mechanism and that NFE2L1 may regulate INSIG1 protein turnover.

To investigate how NFE2L1 affects INSIG1 stability, we expressed V5-tagged INSIG1 with a hepatocyte-specific thyroxine-binding globulin (TBG) promoter in the livers of NFE2L1-WT or - LKO mice using adeno-associated virus (AAV). Three weeks post-viral administration, primary hepatocytes were isolated, and the rate of INSIG1 degradation was monitored following cycloheximide (CHX) treatment. Notably, in the WT hepatocytes, a substantial fraction of INSIG1 disappeared within 60 minutes, reflecting a relatively short half-life of about 30 minutes [20]. By contrast, in the NFE2L1-deficient hepatocytes, INSIG1 levels remained 2-fold higher than the WT after 60 minutes, indicating a significantly extended protein stability **(Fig. 2I-J)**. To determine whether the effect of NFE2L1 on SREBP1 activity is dependent on INSIG1 *in vivo*, we next assessed exogenous INSIG1-V5 levels in the liver. Consistently, liver tissue of NFE2L1-LKO mice exhibited significantly elevated INSIG1 levels compared to WT controls under both fasting and refeeding conditions. Upon refeeding, levels of the SREBP1 were also reduced in the liver tissue from LKO mice (**Fig. S3A**). Taken together, these results suggest that liver NFE2L1-deficiency leads to increased hepatic INSIG1 levels, which in turn reduces SREBP1 activity and ultimately hindering VLDL biogenesis and secretion.

### NFE2L1 interacts with INSIG1 at the ER to facilitate INSIG1 degradation

We next dissect how NFE2L1 regulates INSIG1 stability. NFE2L1, an ER membrane protein, cycles between the ER (as a 130 kDa glycosylated precursor) and nucleus (as a proteolytically cleaved 100 kDa form) depending on proteasome activity [16]. Given their shared ER localization and sterol sensing ability, we hypothesized that NFE2L1 regulates INSIG1 through a direct interaction. To address this possibility, we performed co-immunoprecipitation (Co-IP) in hepa1-6 cells co-expressing hemagglutinin (HA)-tagged NFE2L1 and paramyxovirus SV5 (V5)-tagged INSIG1. We showed that V5-tagged INSIG1 specifically pulled down HA-tagged NFE2L1, with the 130 kDa ER resident form as the predominating protein in the complex (**Fig. 3A**). The amount of nuclear 100 kDa NFE2L1 was minimal in the immunoprecipitates, suggesting that the NFE2L1-INSIG1 interaction occurs predominantly in the ER.

**Figure 3.**
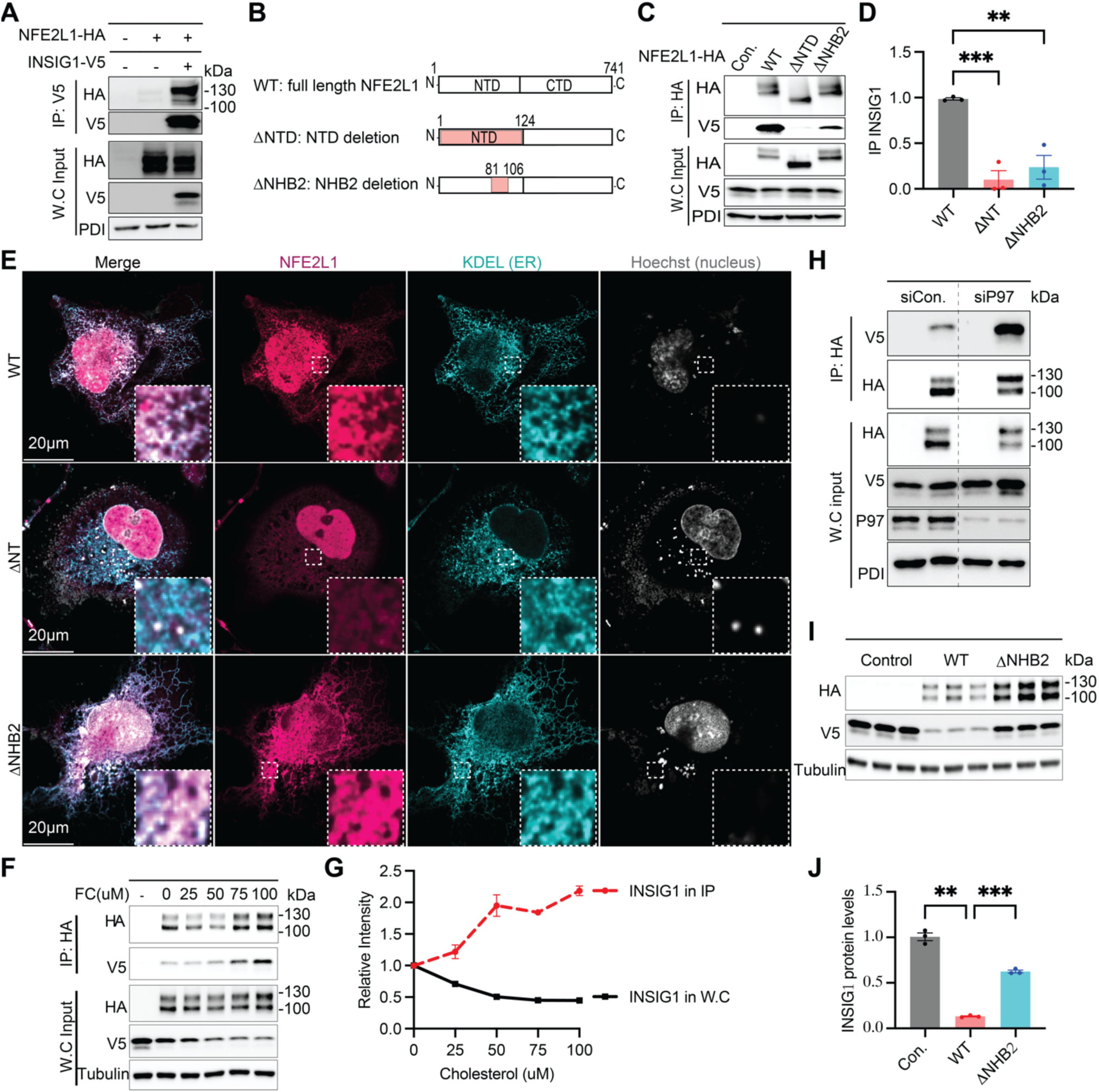
NFE2L1 interacts with INSIG1 in the ER through the NHB2 domain. (**A)** Immunoblot analysis of whole cell lysates and anti-V5 immunoprecipitates from Hepa1-6 cells co-transfected with INSIG1-V5 and NFE2L1-HA constructs. **(B)** Schematic of NFE2L1 domain deletions (NTD and NHB2, red) used for Co-immunoprecipitates (Co-IP) studies. **(C-D)** Co-IP of INSIG1-V5 and HA-tagged NFE2L1-WT or *Δ*NHB2 by anti-HA beads. Quantification of precipitated INSIG1-V5 levels were normalized to NFE2L1-HA. **(E)** Immunofluorescence of COS7 cells expressed NFE2L1-WT, *Δ*NTD and *Δ*NHB2. Cells were stained with Hoechst (nuclear marker) and KDEL (ER marker); images were acquired via confocal microscopy. (**F-G)** Co-IP of INSIG1 and NFE2L1 in Hepa1-6 cells treated with free cholesterol. Quantification of precipitated INSIG1 were normalized to IP NFE2L1. **(H)** Co-IP of INSIG1 and NFE2L1 in Hepa1-6 cells transfected with control or P97 siRNA. **(I-J)** Immunoblot analysis of INSIG1 in hepa1-6 co-transfected with NFE2L1-WT or *Δ*NHB2 and INSIG1-V5. All immunoblots are representative of 3 independent experiments. Data: Means ± SEM. Statistics: One-way ANOVA (**p<0.01, ***p<0.001).

To identify the NFE2L1 domain responsible for INSIG1 binding, we generated truncation mutants (**Fig. 3B, Fig. S3B and D**). Deleting the C-terminal domain (CTD, residues 298-741) had no effect on INSIG1 binding, ruling out CTD involvement (**Fig. S3C)**. We then focused on the N-terminal domain (NTD) and engineered targeted deletions, including ΔNTD (lacking the entire N-terminal domain, residues 1-124, required for ER retention), ΔNHB1 (lacking the N-terminal homology box 1, residues 11-30, also required for ER retention), ΔSAS (lacking a signal peptide-associated sequence, residues 31-50), ΔCRAC1/2 (lacking the putative cholesterol-recognition amino acid consensus sequences, residues 62-82) and ΔNHB2 (lacking the N-terminal homology box 2, residues 81-106) [21, 22] **(Fig. 3B and Fig. S3D).** Deleting the NTD and the NHB2 within the NTD substantially abolished INSIG1 binding (**Fig. 3C-D and Fig. S3E**). The mutant NFE2L1-ΔNTD showed a single band, while the ΔNHB2 protein, like NFE2L1-WT, exhibited two bands: one corresponding to the glycosylated precursor located in the ER and one corresponding to the cleaved form in the nucleus. As shown in previous studies, ΔNTD is predominantly located in the nucleus, whereas ΔNHB2 retains the same subcellular localization and transcriptional activity as NFE2L1-WT [23]. Our confocal microscopy images further confirmed the subcellular location of NFE2L1-WT, -ΔNTD and -ΔNHB2. Like NFE2L1-WT, ΔNHB2 was present in both the ER and nucleus, whereas the majority of ΔNTD was mislocalized to the nucleus **(Fig. 3E)**. Despite its normal ER location, ΔNHB2 exhibited reduced binding to INSIG1, suggesting that the NHB2 domain within the NTD is required for their interaction.

Our previous study demonstrated that free cholesterol retains NFE2L1 in the ER [14]. To tested cholesterol regulation of the NFE2L1-INSIG1 interaction, we treated cells with various cholesterol derivatives: low-density lipoprotein (LDL), free cholesterol (FC), and 25-hydroxycholesterols (25-HC). Notably, only FC increased the amount of INSIG1 associated with NFE2L1 in the precipitates **(Fig. S3F)**. To further validate this finding, we treated cells with increasing levels of cholesterol and observed that the ER-bound NFE2L1 increased progressively with increasing FC levels. Importantly, the amount of INSIG1 pulled down by NFE2L1 was also increased with rising levels of FC (**Fig. 3F-G**). To validate this regulation, we modulated NFE2L1 ER localization by silencing valosin-containing protein (VCP/P97), an ATPase required for NFE2L1 retro-translocation to the cytosol. Knocking down P97 increased amounts of the ER-localized NFE2L1 (130 kDa), along with a corresponding increase in the INSIG1 co-immunoprecipitated by NFE2L1 **(Fig. 3H)**. These results further support that NFE2L1 at the ER senses free cholesterol levels and interacts with INSIG1 at this location.

Interestingly, the interaction inversely correlated with total INSIG1 levels: higher NFE2L1-INSIG1 binding (e.g., under FC treatment) coincided with lower INSIG1 in the whole cell (**Fig. 3F-G and Fig. S3F**). To confirm causality, we expressed NFE2L1-WT, or the reduced INSIG1 binding mutant ΔNHB2 with INSIG1. While NFE2L1-WT reduced INSIG1 protein levels by 90%, the INSIG1-binding-deficient mutant ΔNHB2 exhibited a markedly attenuated effect **(Fig. 3I-J).** This demonstrated that NHB2-mediated binding is essential for INSIG1 degradation. Altogether, these findings suggest that NFE2L1 interacts, via the NHB2 domain, with INSIG1 at the ER and, in response to excessive free cholesterol, to drive INSIG1 degradation.

### The NFE2L1-INSIG1 axis drives SREBP1 activity and VLDL secretion

To establish the physiological relevance of the NFE2L1-INSIG1 interaction, we performed *in vivo* rescue experiments in LKO mice using AAV vectors expressing NFE2L1-WT, or the INSIG1-binding deficient ΔNHB2 mutant. The metabolic parameters of these mice are summarized in **Figure S4A-E**. NFE2L1 expression in the liver did not cause any significant alterations in body weight, food intake, liver weight, or plasma levels of NEFA and insulin.

We fractionated the liver into cytoplasmic and nuclear fractions to compare the protein levels of precursor membrane-bound SREBP1(P) and the active nuclear form SREBP1(N). NFE2L1-LKO mice in the AAV-GFP (LKO_GFP) expressing group exhibited significantly lower levels of nuclear SREBP1(N) compared to WT mice (WT_GFP) (**Fig. 4A-B**). Importantly, reintroducing NFE2L1-WT into LKO livers (LKO_NFE2L1) restored nuclear SREBP1(N) levels to those comparable to WT mice. Conversely, expression of the INSIG1-binding deficient mutant ΔNHB2 (LKO_ΔNHB2) resulted in nuclear SREBP1(N) levels similar to those in the LKO_GFP group, which were significantly lower than those in the WT_GFP and LKO_NFE2L1 mice (**Fig. 4A-B**). Consistently, mRNA expression of SREBP target genes *Hmgcs, Hmgcr, Mvd, Mvk* and *Pcsk9*, were significantly diminished in LKO_GFP mice compared to WT counterparts. Restoration of NFE2L1-WT, but not ΔNHB2 expression in LKO mice, led to the normalization of gene expression (**Fig. S5A-G**). Importantly, the mRNA levels of NFE2L1 target gene *Psmb5* were found to be comparable between the NFE2L1-WT and ΔNHB2 mice (**Fig. S5H),** suggesting that the observed difference in SREBP1 activity are attributable to the interaction between NFE2L1 and INSIG1, rather than differences in transcriptional activity.

**Figure 4.**
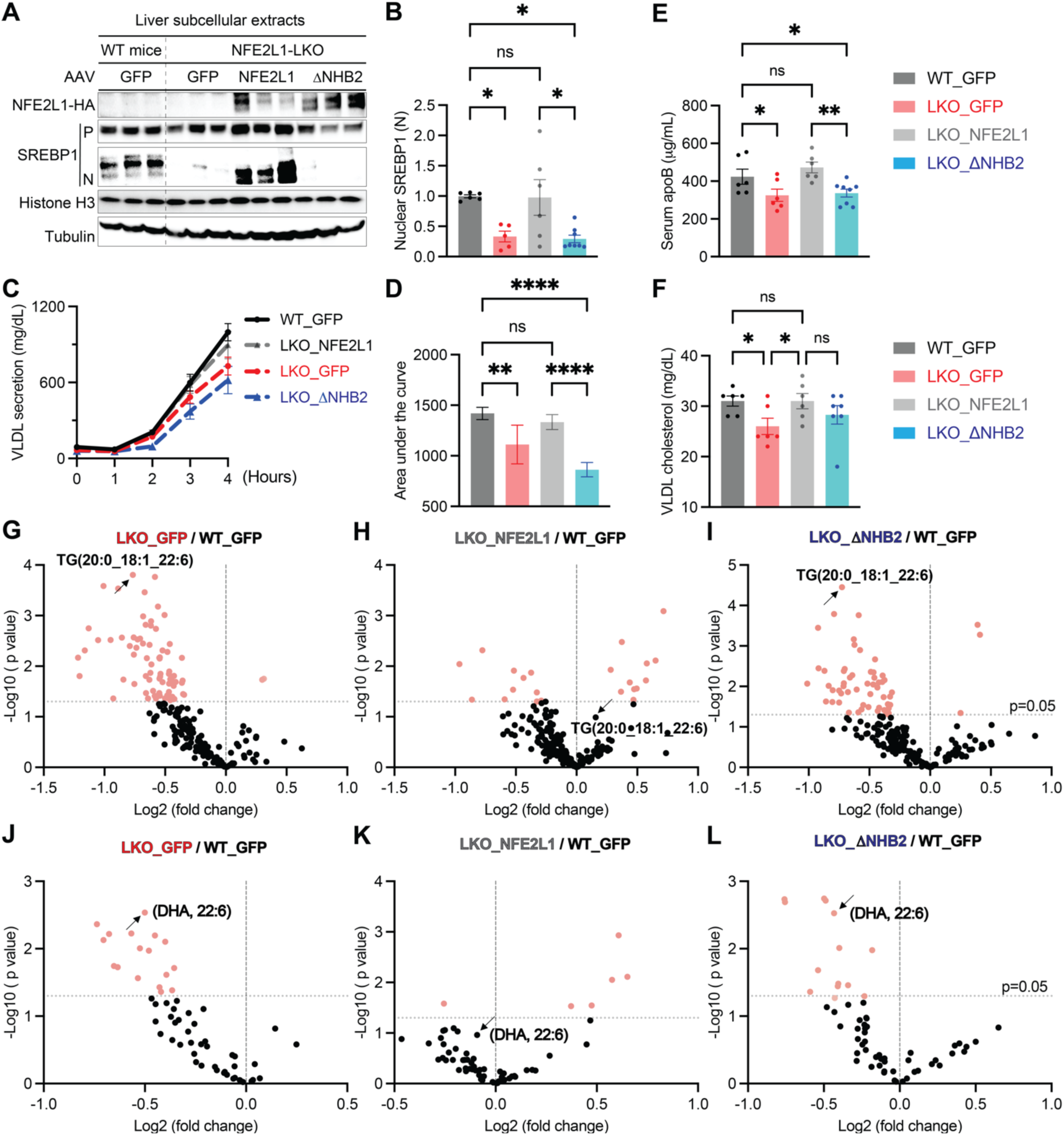
Hepatic NFE2L1-INSIG1 axis regulates SREBP1 and VLDL secretion. 8-week-old NFE2L1-LKO and WT littermates were injected with AAV-GFP, AAV-NFE2L1-WT or AAV-NFE2L1-ΔNHB2. **(A)** Immunoblot analysis of SREBP1 proteins in the liver subcellular extracts. Mice were fasted overnight and refed for 6 hours prior to analysis (n=6-8/group). Tubulin (cytosolic) and histone H3 (nuclear) serve as loading controls. **(B)** Quantification of SREBP1(N) from (A). **(C)** VLDL secretion assay: Mice were fasted overnight, plasma TG levels were measured at 0, 1, 2, 3, and 4 hours post-tyloxapol injection (n=14/group). **(D)** Area under the curve (AUC) for TG secretion in (C). **(E)** Serum apoB lipoprotein levels. **(F)** Serum VLDL cholesterol. **(G-I)** Lipidomic profiling for serum TG species (n=6/group). **(J-L)** Fatty acid composition analysis within serum TG pool (n=6/group). Red denote p<0.05. Data: Means ± SEM. Statistics: *t*-test or One-way ANOVA, *p<0.05, **p<0.01, ****p<0.0001.

To test whether the NFE2L1-INSIG1 axis promotes VLDL secretion *in vivo*, we performed a secretion assay in the LKO mice expressing NFE2L1-WT and NFE2L1-ΔNHB2. Consistent with the reduction of SREBP1 [8], NFE2L1-LKO mice exhibited significantly reduced hepatic VLDL secretion compared to the WT controls (**Figs. 1I-J and 4C-D**). Notably, reintroducing NFE2L1-WT into the LKO mice restores VLDL secretion, whereas expression of the mutant NFE2L1-ΔNHB2 form failed to correct VLDL secretion in the NFE2L1-LKO mice (**Fig. 4C-D**). This observation was further supported by similar patterns detected in serum apolipoprotein B (apoB) and VLDL cholesterol levels (**Fig. 4E-F**). These results underscore the importance of the NFE2L1-INSIG1 axis in modulating VLDL biogenesis and secretion *in vivo*.

To gain further insights into the effect of the NFE2L1-INSIG1 axis on lipid metabolism, we performed lipidomic analysis to determine serum lipid composition in detail. Lipid profiles revealed that NFE2L1 deficiency significantly reduced total TG and diminished most individual TG species, irrespective of their fatty acid composition (**Fig. 4G and Table S1**). Strikingly, while re-expression of NFE2L1-WT restored serum TG abundance and species in LKO mice, the INSIG1-binding-deficient ΔNHB2 mutant failed to rescue this phenotype (**Fig. 4H-I**). For instance, TG (20:0_18:1_22:6), which was markedly reduced in LKO mice compared to WT controls, returned to WT mice levels upon NFE2L1-WT re-expression but remained suppressed in ΔNHB2-expressing mice (**Fig. 4G-I**). This regulatory axis extended to phosphatidylcholine (PC) species, which are essential for VLDL assembly and secretion. PC levels, significantly reduced in LKO mice, were partially rescued by NFE2L1-WT re-expression but not by the ΔNHB2 mutant (**Fig. S6A-C**). These results underscore the necessity of functional NFE2L1-INSIG1 interaction in maintaining hepatic and systemic lipid homeostasis, with the ΔNHB2 mutant’s inability to restore TG, or PC profiles highlighting the critical role of this interaction in coordinating lipogenesis and VLDL secretion.

Parallel lipidomic profiling of the TG pool demonstrated that the NFE2L1-INSIG1 axis selectively modulates polyunsaturated fatty acid (PUFAs) composition. Notably, docosahexaenoic acid (DHA, 22:6), which was depleted in LKO mice, was fully restored by NFE2L1-WT but not by the ΔNHB2 mutant (**Fig. 4J-L and Table S2**). In addition to DHA, we also detected similar patterns for both docosapentaenoic acid (DPA, 22:5), and eicosapentaenoic acid (EPA, 20:5), indicating that anti-inflammatory PUFAs may be preferentially regulated by the NFE2L11-INSIG1 axis. (**Fig. S6D-F**). Interestingly, clinical data show that elevated PUFAs correlate with reduced MASH severity by enhancing VLDL secretion, suppressing liver inflammation, and inhibiting hepatic stellate cells (HSCs) fibrogenesis [24, 25]. These results suggest that modulation of the NFE2L1-INSIG1 axis may counteract important pathways in MASH pathogenesis, a mechanism we directly tested in our *in vivo* model.

### The NFE2L1-INSIG1 axis enhances hepatic cholesterol secretion and attenuate MASH

NFE2L1-deficient mice exhibited liver dysfunction even on a standard chow diet, characterized by 2.5-fold elevated serum alanine transaminase (ALT) and 1.2-fold higher aspartate transaminase (AST) **(Fig. S7A)**, biomarkers indicative of hepatocellular injury. In addition, the LKO mice exhibited a significant upregulation of gene expression associated with liver inflammation (e.g. *Cd68, Ccl2* and *Ccl5)* and fibrosis *(e.g. Tgf-ß, Col1a1* and *Timp-1)* (**Fig. S7B**). These findings highlight the critical role of NFE2L1 in maintaining liver health, consistent with earlier reports [14, 16, 17, 26, 27].

To directly evaluate NFE2L1’s therapeutic potential in MASH, we employed *db/db* mice fed a methionine/choline-deficient (MCD) diet, a severe model mimicking human MASH pathology characterized by impaired VLDL secretion and progressive liver damage [28, 29]. These mice were administrated with adenoviral vectors (AV) expressing either LacZ as a control or NFE2L1 while being fed a MCD diet for two weeks. Overexpression of NFE2L1 robustly activated the SREBP pathway, increasing the hepatic mRNA levels of lipogenic targets **(Fig. 5A**). Immunoblot analysis confirmed the increase of NFE2L1 protein, accompanied by a significant reduction in INSIG1 levels (**Fig. 5B-D**), which facilitated an increase in SREBP1/2 activity. This was further supported by the upregulation of LDL receptor (LDLR) protein (**Fig. 5E**), a key SREBP2 target critical for LDL cholesterol clearance.

**Figure 5.**
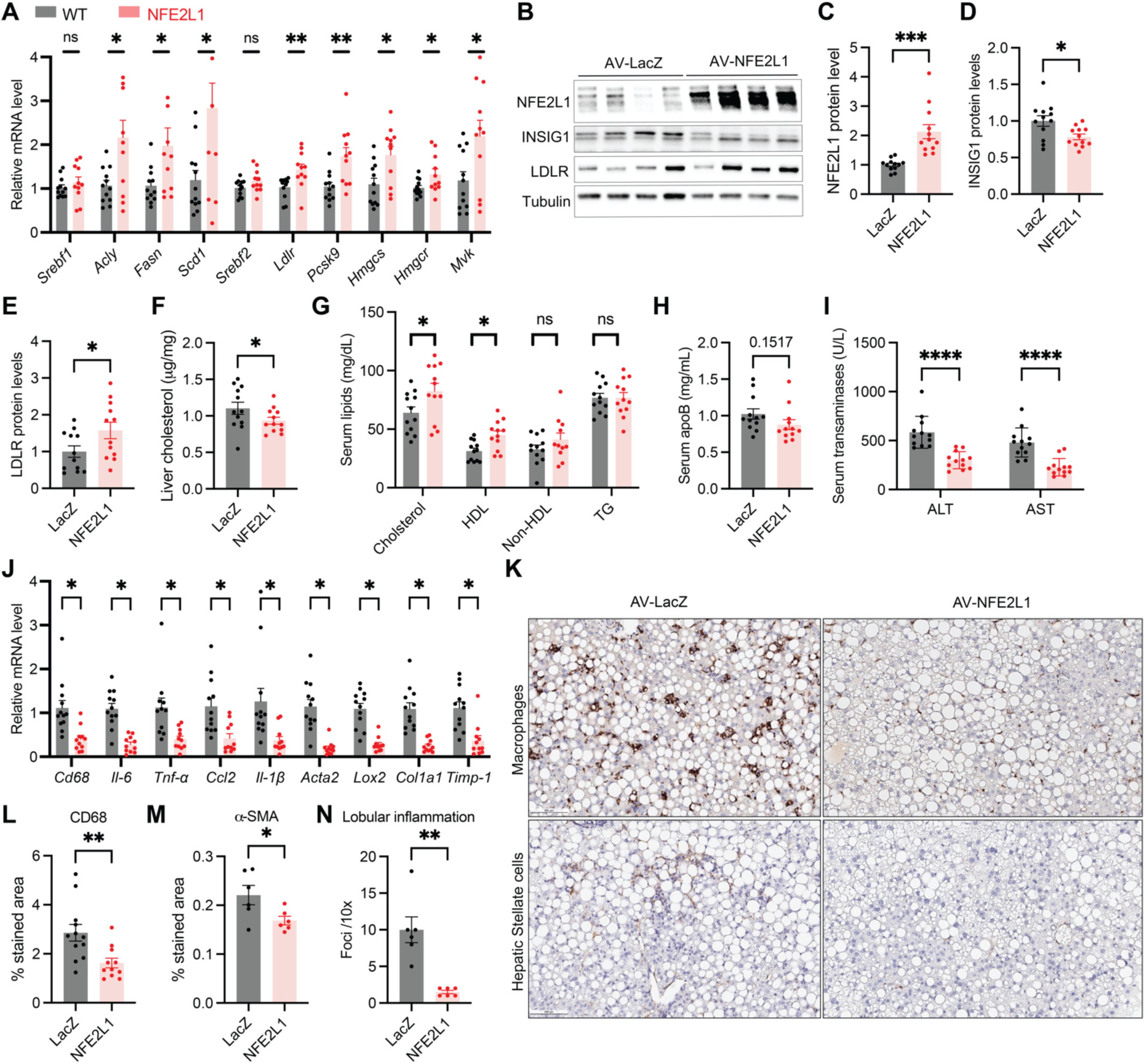
NFE2L1 ameliorates liver injury and inflammation in MASH. 8-week-old db/db mice were injected with either AV-LacZ or AV-NFE2L1 and fed with MCD diet for 2 weeks. Mice were fed ad libitum before samples collection (n=12/group). **(A)** Liver mRNA levels of SREBP1/2 target genes normalized to 18S ribosomal RNA (18S). **(B-E)** Immunoblot and quantification of liver lysates for NFE2L1, INSIG1 and LDLR. **(F)** Liver cholesterol levels. **(G)** Serum lipids profiles measured via Piccolo Lipid Panel Plus. **(H)** Serum apoB level by ELISA **(I)** Serum ALT and AST levels(Piccolo Lipid Panel Plus). **(J)** mRNA levels of liver inflammation and fibrosis markers. **(K-N)** Representative histology images and quantification of liver section staining macrophage marker CD68 and hepatic stellate cell marker *α-*SMA, and lobular inflammation. Data: Mean ± SEM. Statistics: *t*-test (*p<0.05, **p<0.01, ***p<0.001 ****p<0.0001).

Notably, NFE2L1-expressing mice displayed a reduction in liver cholesterol levels, but elevated serum total cholesterol levels compared to the control (**Fig. 5F-G**). This indicated enhanced VLDL-mediated export, suggesting that NFE2L1 overexpression shifted cholesterol flux from hepatic retention to secretion. Importantly, this serum cholesterol increase was redistributed into anti-atherogenic high density lipoprotein (HDL), while atherogenic risk markers such as non-HDL cholesterol (LDL/chylomicron remnants) and apolipoprotein B (ApoB) remained unchanged (**Fig. 5G-H**), likely due to compensatory LDLR-mediated clearance of atherogenic particles.

The metabolic benefits of NFE2L1 extended to ameliorating MASH-associated pathology. Notably, these mice displayed a marked reduction in serum ALT and AST levels, suggesting that NFE2L1 expression ameliorated hepatic injury (**Fig. 5I**). Hepatic expression of inflammation and fibrosis markers, including *Cd68, Il-6, Tnf-α, Ccl2, Il-1β, Acta2, Lox2, Col1a1, and Timp-1*, were significantly decreased in the livers of NFE2L1-expressing mice compared to the controls (**Fig. 5J**), consistent with attenuation of inflammation and stellate cell activation. Histopathological analysis corroborated these findings: CD68^+^ macrophage infiltration declined by 45%, and α-SMA^+^ activated stellate cells diminished by 24% in the tissue upon NFE2L1 expression (**Fig. 5K-M**). These histological observations, in addition to decreased lobular inflammation (**Fig. 5N**), were independently confirmed by two pathologists blindly, further validating that NFE2L1 overexpression leads to a significant reduction in inflammation and fibrosis in this model of MASH. Our results indicate that NFE2L1 reduces macrophage infiltration to the liver and decreases the activation of hepatic stellate cells (HSCs), which are the hallmarks of liver inflammation and fibrosis. These observations suggest potential pathways for targeting NFE2L1 to mitigate MASH pathogenesis.

## DISCUSSION

Our findings reveal a novel mechanism by which NFE2L1 maintains hepatic lipid and cholesterol homeostasis through its interaction with INSIG1. We demonstrate that NFE2L1 deficiency elevates INSIG1 levels, suppressing SREBP1 activation and impairing VLDL secretion, which drives hepatic cholesterol accumulation and spontaneous liver injury. Conversely, NFE2L1 overexpression in *db/db* mice fed a methionine-choline-deficient diet restores SREBP1 activity, enhances VLDL-mediated lipid export, and attenuates MASH hallmarks - liver injury, inflammation, and fibrosis. Central to this regulatory axis is the NHB2 domain of NFE2L1, which directly binds INSIG1 in the ER, destabilizing it to enable SREBP1 processing and activation. Rescue experiments confirm the necessity of this interaction: NFE2L1-WT restores lipid homeostasis in knockout mice, while the ΔNHB2 mutant (INSIG1 binding-deficient) fails to rescue SREBP1 activation or VLDL secretion. Lipidomic profiling further underscores the specificity of this axis, as NFE2L1-WT, but not ΔNHB2, reverses alterations in serum triglycerides and fatty acid composition, notably restoring anti-inflammatory PUFAs such as DHA and EPA.

The interaction between NFE2L1 and INSIG1 is dynamically modulated by free cholesterol levels (Fig. 3F-G), enabling NFE2L1 to calibrate its interaction with SREBP based on ER lipid status. This creates a feedback loop, in which NFE2L1 licenses SREBP activation under cholesterol-replete conditions while simultaneously coupling it to VLDL-mediated cholesterol export, a “lipid release valve” mechanism that prevents lipotoxicity by balancing lipogenesis with secretion. Such coordination reconciles the dual roles of SREBP in lipid metabolism: while chronic overactivation is often linked to steatosis, its downregulation impairs VLDL secretion and exacerbates MASH [30, 31], underscoring the necessity of balanced SREBP activity. Our work aligns with evidence that context-specific SREBP activation can optimize lipid dynamics. For example, INSIG1 suppression in mice enhances SREBP activity but reduces hepatic lipid retention by coupling lipogenesis to secretion and improve liver function and reduce inflammation [32]. Similarly, statins, which indirectly activate SREBP2, improve MASH outcomes in humans [33, 34], mirroring our observation that NFE2L1-driven SREBP activation alleviates liver disease.

NFE2L1 overexpression in a *db/db* mice reduced hepatic cholesterol while increasing serum cholesterol via enhanced VLDL secretion, attenuating inflammation and fibrosis. Importantly, this intervention did not elevate atherogenic ApoB or non-HDL levels, due to compensatory LDLR upregulation, a safety feature mirroring statin effects. These finding may suggest potential translation against MASH, a disease affecting 3-5% of the global population with limited treatments [35, 36]. However, broader applicability of the *in vivo* function of NFE2L1 must be validated in diverse models, such as high-fat high-cholesterol diet or toxin-induced MASH.

While NFE2L1 is established as a transcription factor regulating proteasome expression and antioxidative responses [37, 38], our work here uncovers a non-transcriptional role for its ER resident form in sensing cholesterol to promote lipid secretion. By balancing lipogenesis with VLDL-driven secretion, NFE2L1 ensures lipid demands are met without compromising cellular resilience. NFE2L1’s preferential elevation of HDL-associated cholesterol and anti-inflammatory PUFAs suggests a favorable cardiovascular safety profile associated with the mechanism described in this study. This aligns with prior studies showing that SREBP1-mediated lipogenesis supplies essential substrates essential for VLDL assembly [5, 10], yet underscores the need for rigorous cardiovascular risk assessment in future studies. Testing NFE2L1 activation in atherosclerosis-prone models could further clarify its impact on vascular health while reinforcing its role in hepatic protection. Several unanswered questions remain. The detailed mechanism underlying NFE2L1-mediated INSIG1 degradation, including the identification of the responsible E3 ligase and regulatory cofactors, is currently not unknown and warrants investigation. It will also be interesting to explore hormonal or dietary modulation of this axis (e.g., insulin/glucagon, fasting-refeeding cycles) and the long-term consequences of NFE2L1 activation.

Our work defines NFE2L1 as a guardian of hepatic metabolic flexibility, orchestrating SREBP1 activity through INSIG1 destabilization to balance lipid synthesis with secretion. By resolving the paradox of SREBP’s dual roles and the interactions with NFE2L1, our findings advance a mechanism where coordinated activation of lipid pathways mitigate MASH without exacerbating systemic risk. Future efforts to modulate the NFE2L1-INSIG1 axis may offer strategies for metabolic liver disease, addressing a critical unmet need in global health.

## RESOURCE AVAILABILITY

### Lead contact

Further information and requests for resources and reagents should be directed to and will be fulfilled by the lead contact, Gökhan S. Hotamisligil.

### Materials availability

All materials used in the analysis are available to any researcher for purposes of reproducing the analysis.

### Data and code availability

- All data deported in this paper will be shared by the lead contact upon request.
- Original western blot images have been deposited at Mendeley and are publicly available as of the date of publication. Microscopy data reported in this paper will be shared by the lead contact upon request.
- All original code has been deposited at Zenodo and is publicly available as of the date of publication.
- Any additional information required to reanalyze the data reported in this paper is available from the lead contact upon request.

## ACKNOWLEDGMENTS

We thank the Hotamisligil lab members for their support and valuable discussions. We also extend our gratitude to Drs. Scott Widenmaier and Benjamin Garfinkel for their invaluable guidance and maintenance of the mice models, and to Dr. Kacey Prentice, now at the University of Toronto, for her insightful comments on the manuscript. Special thanks to Dr. Renata Goncalves, Zeqiu Wang and Jillian Riveros for their technical assistance with the VLDL secretion assay and primary hepatocyte isolation. We thank Dr. Alba Diaz at the Hospital Clinic of Barcelona for reviewing liver slides from the *db/db* model and quantifying lobular inflammation and steatosis. I.G was funded by a grant from the Instituto de Salud Carlos III (PI22/00776 and BA20/00016).

## AUTHOR CONTRIBUTIONS

S.D. designed the study, performed experiments, analyzed data, interpreted results, and wrote the manuscript. G.Y.L., J.F., G.A., Z.C., I.G., B.Y., and K.E.I. performed experiments and provided expertise and critical feedback. G.S.H. designed and supervised the study, interpreted results, generated project resources, and wrote the manuscript together with S.D. All authors reviewed and commented on the manuscript.

## DECLARATION OF INTERESTS

The authors declare no competing interests.

### DECLARATION OF GENERATIVE AI and AI-ASSISTED TECHNOLOGIES

During the preparation of this work, the authors used ChatGPT to improve the text. After using the tool, the authors reviewed and edited the content as needed and take full responsibility for the content of the publication.

## SUPPLEMENTAL INFORMATION

Supplemental information includes seven figures and two tables and can be found with this article online.

## STAR★METHODS

### KEY RESOURCES TABLE

| REAGENT or RESOURCE | SOURCE | IDENTIFIER |
| --- | --- | --- |
| **Antibodies** |  |  |
| Anti-Histone H3 | Cell Signaling Technology | Cat# 9715 |
| Anti-SCAP | Abcam | Cat# ab153933 |
| Anti-NFE2L1 | Cell Signaling Technology | Cat# 8052; RRID: AB_11178947 |
| Anti-PDI | Cell Signaling Technology | Cat# 3501; RRID: AB_2156433 |
| Anti-V5 | Thermo scientific | Cat# R961-25 |
| Anti-HA | Cell Signaling Technology | Cat# 3724 |
| Anti-AKT | Cell Signaling Technology | Cat# 4691 |
| Anti-pAKT | Cell Signaling Technology | Cat# 9271 |
| Anti-INSIG1 | Abcam | Cat# ab70784 |
| Anti-SREBP1 | Abcam | Cat# ab28481 |
| Anti-CD68 | Proteintech | Cat# 25747-1-AP |
| Anti-LDLR | Abcam | Cat# ab52818 |
| Anti-beta Tubulin HRP | Cell Signaling Technology | Cat# ab21058 |
| **Chemicals, Peptides, and Recombinant Proteins** |  |  |
| Bortezomib (PS-341) | Selleckchem | Cat# S1013 |
| Epoxomicin | Millipore | Cat# 324800 |
| Methyl-β-cyclodextrin | Sigma Aldrich | Cat# C4555 |
| TRIzol | ThermoFisher Scientific | Cat# 15596018 |
| Cholesterol | Sigma Aldrich | Cat# C8667 |
| 25-hydroxycholesterol | Sigma Aldrich | Cat# H1015 |
| Nitrocellulose membrane | Bio-Rad | Cat# 162-0115 |
| Dulbecco's modified medium (DMEM) | GIBCO | Cat# 11965 |
| Cosmic Calf Serum (CCS) | Hyclone | Cat# SH30087.03 |
| Sandoz 58-035 (ACAT inhibitor) | Sigma-Aldrich | Cat# S9318-5MG |
| Fos-Choline-13 | Anagrade | Cat# 85775-42-4 |
| Methyl-β-cyclodextrin | Sigma Aldrich | Cat# C4555 |
| Antibiotics | Lonza | Cat# 17-603E |
| **Bacterial and Virus Strains** |  |  |
| Non-replicating adenovirus expressing β-galactosidase (AV-LacZ) | Vector Biolabs | Cat# 1080 |
| Non-replicating adenovirus expressing wild type mouse Nfe2l1 (AV-NFE2L1) | Vector Biolabs | N/A |
| **Critical Commercial Assays** |  |  |
| Total Cholesterol Assay Kits | Cell Biolabs Inc. | Cat# STA-384 |
| Piccolo Lipid Panel Plus | Abaxis | Cat# 07-P02-12A |
| Nuclear/cytoplasmic fractionation kit | Ambion | Cat# AM1921 |
| Cholesterol Quantitation Kit | Sigma Aldrich | Cat# MAK043 |
| NE-PER™ Nuclear and Cytoplasmic Extraction Reagents | Thermo Scientific | Cat# 78833 |
| Experimental Models: Cell Lines and Mice |  |  |
| Hepa1-6 cells | ATCC | CRL-1830 |
| Primary murine hepatocytes | This Paper | N/A |
| *db/db* mice | The Jackson Laboratory | Jax: 000664 |
| Nfe2l1-Alb-Cre mouse | The Jackson Laboratory | Jax: 003574 |
| Software and Algorithms |  |  |
| Prism 10.0 | GraphPad Software | https://www.graphpad.com |
| ImageJ | NIH | https://imagej.nih.gov/ij/ |
| BioRender |  | https://biorender.com |

### METHOD DETAILS

#### Cell Lines and Cell Culture Conditions

Murine Hepa1-6 cells were cultured in DMEM (Dulbecco’s Modified Eagle Medium) supplemented with 5% Cosmic Calf Serum (CCS). Other cell culture media conditions are described below. All cells were cultured at 37°C in a humidified incubator maintained at a CO2 level of 10%. For gene silencing experiments, Hepa1-6 cells were transfected with specific siRNAs targeting NFE2L1, SCAP or INSIG1 using Lipofectamine RNAiMAX (Thermo Fisher) according to the manufacturer’s instructions. A control siRNA (non-targeting) was used as a negative control. Briefly, 1 × 10^5^ cells were seeded in 6-well plates 24 hours prior to transfection. The siRNA and Lipofectamine RNAiMAX reagent were incubated in Opti-MEM medium (Thermo Fisher) for 15 minutes before being added to the cells. After 48 hours of transfection, cells were harvested for further analyses including Western blotting, quantitative PCR, and co-immunoprecipitation (Co-IP) experiments. For primary mouse hepatocytes isolation, mice were anesthetized with 100 mg/kg xylazine, and the liver was perfused with warm PBS containing 0.5 mM EGTA, followed by digestion with a collagenase solution (collagenase type IV, Sigma) for 15-20 minutes at 37°C. The digested liver was then minced and filtered through a 100 µm cell strainer to obtain a single-cell suspension. Hepatocytes were plated in collagen-coated culture plates at a density of 1 × 10^6^ cells per well in Williams’ E medium (Thermo Fisher) supplemented with 10% FBS and 1% penicillin-streptomycin. Cells were cultured at 37°C in a humidified incubator at 5% CO2. Hepatocytes were used within 24 hours of isolation.

#### Cloning and Mutagenesis to Generate Mutant forms of NFE2L1

To generate the NFE2L1 mutants, site-directed mutagenesis was performed using the Q5 Site-Directed Mutagenesis Kit (New England Biolabs) according to the manufacturer’s instructions. Briefly, the full-length NFE2L1 cDNA was cloned into the pcDNA5.0 vector using standard cloning methods. The NFE2L1-WT construct was used as the template for generating the mutants. For NFE2L1-ΔNHB2, a specific mutation was introduced to delete the NHB2 domain (amino acids 81–106) by introducing a stop codon. This was achieved using the forward primer (CGAGACCCGGAGGGGTCT) and the reverse primer (CCGGGCAGTGAAGTAATTGTCC). The mutated construct was verified by Sanger sequencing to confirm the deletion of the NHB2 domain. The NFE2L1-WT and NFE2L1-ΔNHB2 plasmids were subsequently used for transfection into Hepa1-6 cells for functional analysis, including protein expression and co-immunoprecipitation.

#### Immunoprecipitation and Immunoblotting Experiments

Tissues and cells used for immunoblotting experiments were lysed in buffer containing 150 mM NaCl, 50 mM Tris-HCl (pH 7.4), 1% v/v Nonidet P-40, 5 mM EDTA and fresh protease inhibitors. Lysate was incubated on ice for 30 minutes and then centrifuged at 4°C for 15 min at 13,000 g and the supernatant was transferred to a fresh tube. Protein concentrations were determined using the Pierce BCA Protein Assay kit from ThermoFisher Scientific, and samples were stored in Laemmli buffer at -20°C until analysis. Electrophoresis of protein was run at neutral pH in 4– 20% Criterion™ TGX Stain-Free™ Protein Gel (Bio Rad). Protein in gels were transferred to nitrocellulose membrane, blocked in TBS-T containing 1% milk, and then incubated overnight with primary antibodies in TBS-T containing 1% milk at 4°C with shaking. After washing, anti-rabbit or anti-mouse HRP-conjugated antibodies were applied for 30 min at room temperature. Membranes were incubated with Super Signal West Femto (Thermo Scientific) to generate a chemiluminescent signal and images were captured using Image Lab software (Bio-Rad).

#### Cholesterol Treatments in Cultured Cells

We first solubilized dry cholesterol powder in 100% ethanol at 65°C and then added this to methyl-b-cyclodextrin (MbCD) in water at a concentration of 5 mM cholesterol to 42 mg/ml MbCD, which was then cooled and sterile filtered. Cells were treated by adding the cholesterol complex to media at a concentration of 50 mM (or otherwise at indicated doses) for 3 hours. This method causes cells to accumulate cholesterol because the stoichiometric capacity of MbCD to complex with cholesterol is near maximal. As such, in cell culture media there is a high tendency for cholesterol to transfer into cellular membranes, at which point it circulates through the normal cellular cholesterol pools.

#### Immunofluorescence Analysis

COS-7 cells transiently transfected with wild-type or mutant variants of HA tagged NFE2L1 were seeded on poly-L-lysine coated coverslips. After 2-hour treatment with indicated chemicals, cells were fixed with 3% paraformaldehyde for 15 min, washed 2x in PBS, permeabilized with 0.1% Triton X-100 in PBS (Triton/PBS) for 15 min, and then washed 2x in PBS. Antibodies were diluted in blocking buffer plus 0.05% Triton X-100. Primary antibody immunodetection was performed overnight at 4°C, followed by 2x Triton/PBS washes. Secondary antibody incubation was performed at room temperature for 1 hr, followed by 2x Triton/PBS washes and 2x PBS washes. Preparations were further incubated with DAPI in PBS (1/10,000) for 15 min, followed by 2 additional PBS washes. Images were taken in our imaging lab in a Leica SP8 X confocal microscope equipped with 405 nm and White Light (WLL) lasers and a 63x oil immersion objective.

#### Subcellular Extraction of Liver Tissues

Liver tissues from mice were processed for subcellular fractionation using Thermo Scientific NE-PER Nuclear and Cytoplasmic Extraction system following the manufacturer’s instructions. Briefly, livers were harvested and immediately placed in cold PBS. The tissue was then homogenized in the cytoplasmic extraction buffer, supplemented with protease inhibitors. The homogenates were centrifuged at 1,000 × g for 10 minutes at 4°C to remove debris, and the supernatant was collected as the cytoplasmic fraction. For nuclear extraction, the remaining pellet was resuspended in the nuclear extraction buffer, incubated on ice for 30 minutes, and then centrifuged at 14,000 × g for 10 minutes at 4°C to separate the nuclear fraction. The supernatant was discarded, and the nuclear pellet was resuspended for subsequent analysis. Both the nuclear and cytoplasmic fractions were stored at -80°C until used for protein quantification and Western blot analysis. Protein concentrations in the fractions were determined using BCA protein assay kit, and equivalent amounts of protein from both fractions were used for analysis of target proteins by Western blotting.

#### Quantitative PCR

Cells or tissues were homogenized in TRIzol Reagent for total RNA isolation. Complementary DNA (cDNA) was synthesized iScript™ cDNA Synthesis Kit. The qPCR was performed on a ViiA7 system (Applied Biosystems) using SYBR green. Gene of interest cycle thresholds (Cts) were normalized to 18S levels by the DDCt method and displayed as expression levels relative to controls.

#### Mouse Studies

All mice were bred and housed on a 12 hr light/dark cycle in the Harvard T.H. Chan School of Public Health pathogen-free barrier facility with ad libitum access to standard chow diet (5053 - PicoLab Mouse Diet from LabDiet), unless otherwise specified. Males were used for all experiments and were performed at the age of 8-10 weeks. Liver-specific Nfe2l1 deficient (NFE2L1-LKO) mice were generated by Nfe2l1-floxed mice crossed with hemizygous B6. Cg-Tg (Alb cre) 21Mgn/J transgenic mice expressing Albumin promoter driven Cre recombinase. Genotyping for Nfe2l1 flox and Cre recombinase were performed by real time PCR with specific probes designed for each gene (Transnetyx, Cordova, TN). Mice were injected intravenously via retro-orbital plexus with 1×10^11^ viral genomes of AAV-NFE2L1-WT, AAV-NFE2L1-ΔNHB2, or AAV-GFP. Three weeks after injection, mice were subjected to fasting (16 hours) followed by refeeding (6 hours). Mice were then euthanized, and tissues and serum were collected for further analysis. 8-week-old male *db/db* mice were placed under isoflurane anesthesia (1-3%) and dosed intravenously via the retro-orbital plexus with 1 x 10^9^ particles/mouse (Ad-LacZ, Ad-NFE2L1, Vector Biolabs) on day 0. On day 2 after virus injection mice were started on MCD diet. Mice were euthanized on day 14. All mouse studies were approved by the Harvard Medical Area Standing Committee on Animals.

#### Serum NEFA, Triglyceride and Insulin Measurements

Blood samples were collected via cardiac puncture from euthanized mice or from tail vein of liver mice and were centrifuged at 1500 x g for 10 minutes to obtain serum. Serum non-esterified fatty acid (NEFA) levels were measured by enzymatic assay using a commercially available NEFA measurement system (Wako Chemicals, Richmond, VA). Serum triglyceride (TG) levels were quantified using the Triglyceride Quantification system (Thermo Fisher, Catalog #TR0100). The TG concentration in each serum sample was determined from a standard curve generated with known concentrations of TG standards. Serum insulin levels were quantified using an enzyme-linked immunosorbent assay (ELISA) according to the manufacturer’s instructions (Mouse Insulin ELISA, Crystal Chem, Downers Grove, IL).

#### Liver and Serum Cholesterol Measurement

Cholesterol levels in liver tissue and serum were measured using the Cholesterol Quantitation system (Sigma Aldrich, Cat# MAK043). To quantify cholesterol levels, 10 µL of serum was diluted with 90 µL of the provided cholesterol assay buffer. The cholesterol levels were measured by adding the working reagent, which contains cholesterol esterase, cholesterol oxidase, and peroxidase, to each sample and incubating for 30 minutes at 37°C. The absorbance was measured at 570 nm, and cholesterol concentrations were calculated from a standard curve generated with known cholesterol standards. Liver tissue was homogenized in the provided cholesterol assay buffer. Briefly, 100 mg of liver tissue was homogenized in 1 mL of buffer, and the homogenate was centrifuged at 10,000 × g for 10 minutes at 4°C to remove debris. The supernatant was collected, and protein concentration was determined using the BCA protein assay. For cholesterol quantification, 20 µL of liver supernatant was mixed with 180 µL of cholesterol assay buffer, followed by the addition of the working reagent. The reaction was incubated at 37°C for 30 minutes, and absorbance was measured at 570 nm.

#### Serum and Liver Lipidomic

Lipidomics were performed by the Harvard Chan Advanced Multi-Omics Platform, directed by Dr. Sheng (Tony) Hui. Lipids were extracted in butanol/methanol (1:1) with 5 mM ammonium formate. 20 µL extraction solvent was used per mg tissue, or 10 µL per µL serum. The mixture was vortexed vigorously and then centrifuged at 13,000 × g for 10 minutes at 4°C. 30 μL of the supernatant was transferred to an HPLC vial with glass insert, and 5 μL of this sample was injected into the liquid chromatography-mass spectrometry (LC-MS) system for analysis. The instrument used was Dionex Ultimate 3000 - QExactive mass spectrometry (Thermo Scientific, Waltham, MA, USA). Chromatographic separation was achieved on an Acquity UPLC CSH C18 column (130Å, 1.7 μm, 2.1 mm × 100 mm) at 50°C. Mobile phase A was acetonitrile and water (6:4), and phase B was isopropanol and acetonitrile (9:1), both phases containing 10 mM ammonium formate and 0.1% formic acid. The elution gradient was 0-3 min, 20% B; 3-7 min, 20-55% B; 7-15 min, 55-65% B; 15-21 min, 65-70% B; 21-24 min, 70-100% B; and 24-26 min, 100% B, 26-28 min, 100-20% B, 28-30 min, 20% B, with a flow rate of 0.35 mL/ min. The autosampler was at 4°C. The injection volume was 5 μL. MS analysis was performed in positive and negative ionization polarities using a combined full mass scan and data-dependent MS/MS (Top 10) (Full MS/dd-MS2) approach. Precursor ion scan had resolving power 70,000, covering 100-1200 *m/z*. For product ion scan, resolving power was 17,500, isolation width 1 *m/z*, and stepped normalized collision energy 10, 20, and 40 eV. The intensity threshold of precursor ions for dd-MS2 analysis and the dynamic exclusion were set to 1.6 × 10^5^ and 10 s, respectively. Thermo Scientific LipidSearch software version 5.0 was used for lipid identification and quantitation. First, the product search mode was used to identify lipids based on the exact mass of the precursor ions and the MS2 mass spectra of product ion scan. The precursor and product tolerance were 10 ppm. The absolute intensity threshold of precursor ions and the relative intensity threshold of product ions were set to 30000 and 1%, respectively. Next, the search results from all samples were aligned within a retention time tolerance of 0.25 min.

#### In Vivo VLDL secretion

Male were fasted for 16 hours (3 pm-7 am). A baseline (0 h) blood sample was then collected from the tail vein followed by an intraperitoneal injection of 10% Tyloxapol/saline solution (Sigma-Aldrich) (500 mg/kg body weight). Blood was then collected from the tail vein at 1, 2, 3 and 4 hours after the injection and plasma was separated for measurement of TG levels. The plasma TG secretion rate was calculated from the slope of the linear regression of the time vs. TG concentration.

#### Histology

Livers were fixed in 10% zinc formalin overnight and then transferred to 70% ethanol for further preservation. Tissue processing, sectioning, and staining with hematoxylin and eosin (H&E) were performed by HistoWiz, a fee for service histology service provider. The stained tissue sections were used for assessment. Histopathological evaluations were conducted by two independent pathologists, included quantification of lobular (foci) inflammation.

### QUANTIFICATION AND STATISTICAL ANALYSIS

Statistical significance was assessed using GraphPad Prism 10. For comparisons between two groups, significance was determined using a two-tailed unpaired t-test. Multiple group comparisons were analyzed by one-way analysis of variance (ANOVA) was performed. For comparisons between groups over multiple time points two-way ANOVA were performed. A p-value threshold of < 0.05 was considered statistically significant. Unless otherwise stated, all data are presented as the mean ± standard error of the mean (SEM).

## SUPPLEMENTAL FIGURES

**Figure S1.**
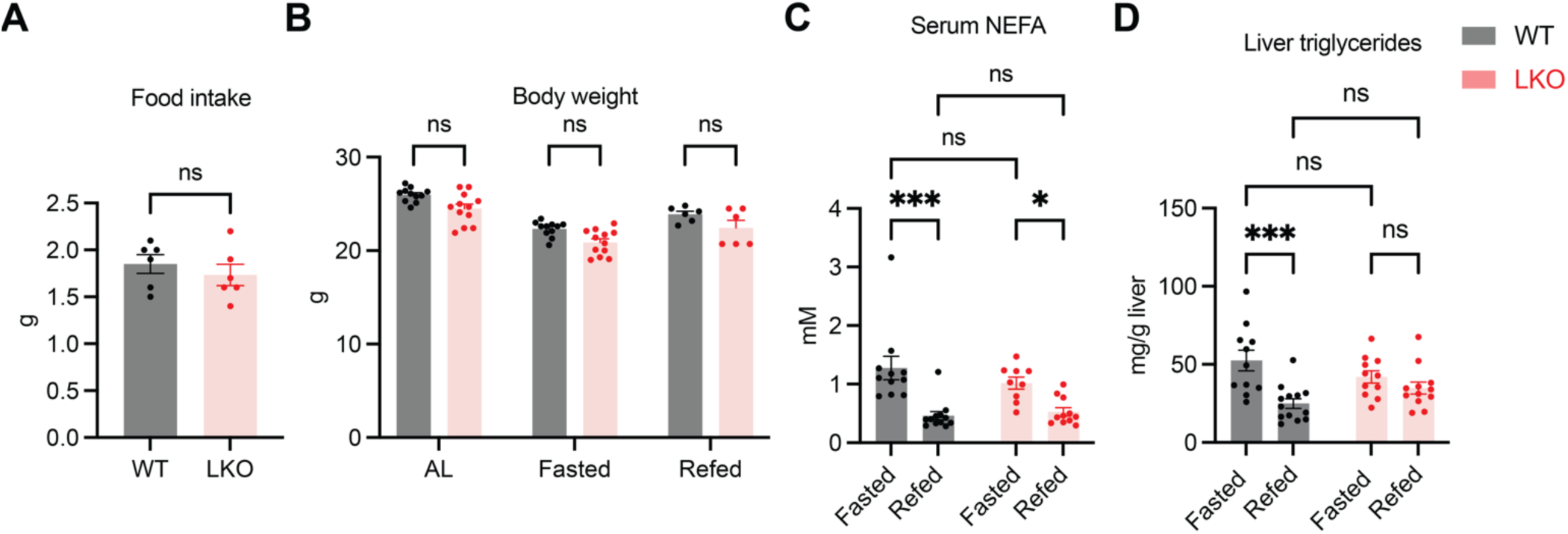
NFE2L1 deficiency does not alter food intake, body weight, or serum NEFA. **(A)** Food intake during a 6-hour refeeding period after 16-hour fasting in NFE2L1 WT and LKO mice (n=6/group). **(B)** Body weight of WT and LKO mice under ad libitum (AL), 16-hour fasted (Fasted), and 6-hour refed (Refed) conditions. **(C)** Serum NEFA levels measured by ELISA (n=11/group). (D) Liver TG levels. Data are mean ± SEM. Statistical significance assessed by two-way ANOVA or Student’s t-test; *p<0.05, ***p<0.001.

**Figure S2.**
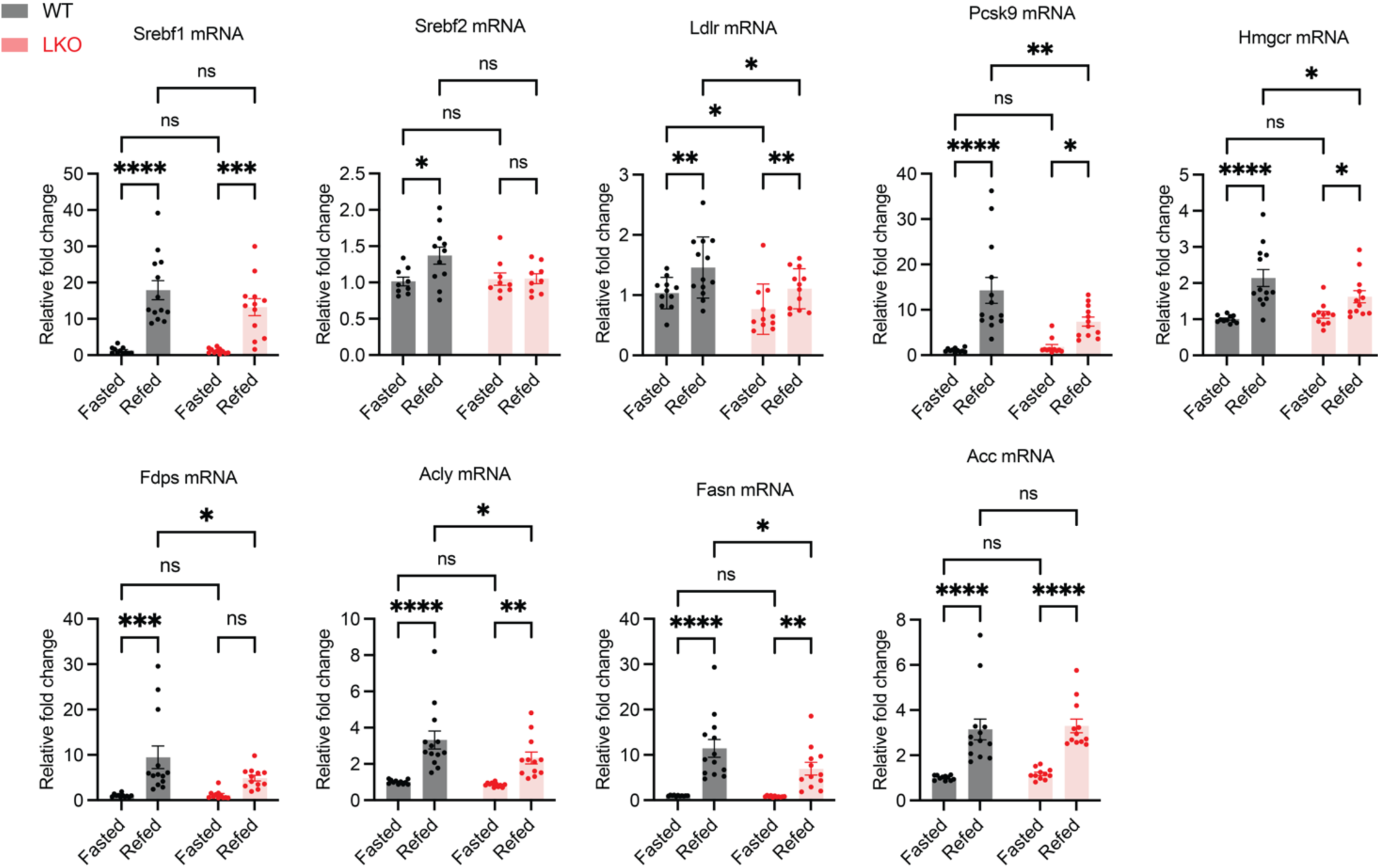
Hepatic mRNA levels of SREBP target genes in WT and LKO mice. 8-week-old NFE2L1 WT or LKO mice were fasted for 16 hours or fasted for 16 hours followed by 6-hour refeeding (n=11/group). Gene expression was normalized to 18S ribosomal RNA. Gray and red bars denote WT and LKO mice, respectively. Statistical significance was assessed by two-way ANOVA; P < 0.05, *P < 0.01, **P < 0.001, ***P < 0.0001.

**Figure S3.**
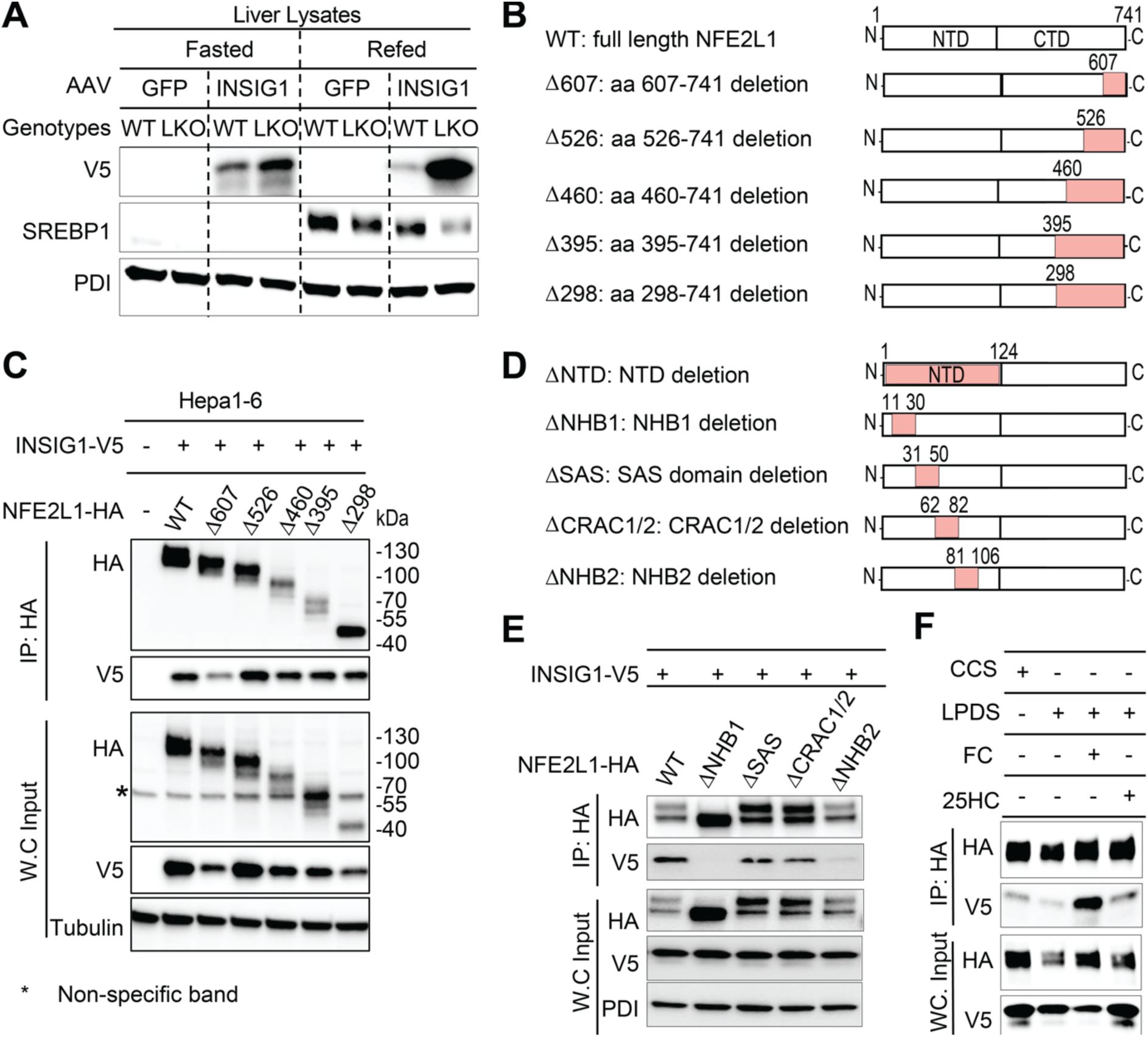
The N terminal domain (NTD) of NFE2L1 binds to INSIG1 in the ER. **(A)** Immunoblotting analysis of INSIG1-V5 in liver from WT and LKO mice injected with AAV-GFP or INSIG1-V5 under fasting or refed conditions. **(B)** Schematic of NFE2L1 truncation mutants lacking regions in the C-terminal domain (CTD). Red indicates deleted regions. **(C)** Co-immunoprecipitation (Co-IP) of INSIG1-V5 with HA-tagged NFE2L1 CTD mutants in Hepa1-6 cells. Lysates were harvested 48 h post-transfection and immunoprecipitated with HA beads. **(D)** Schematic of NFE2L1 truncation mutants lacking regions in the N-terminal domain (NTD). **(E)** Co-IP of INSIG1-V5 with HA-tagged NFE2L1 NTD mutants, performed as in (B). **(F)** Sterol-dependent interaction between NFE2L1 and INSIG1. Cells transfected with NFE2L1-WT and INSIG1-V5 were treated for 3 h with: 5% cosmic calf serum (CCS), 5% lipoprotein-deficient serum (LPDS), 50 µM free cholesterol (FC), or 1 µM 25-hydroxycholesterol (25HC). Lysates were subjected to Co-IP with HA beads. Immunoblots are representative of ≥3 independent experiments.

**Figure S4.**
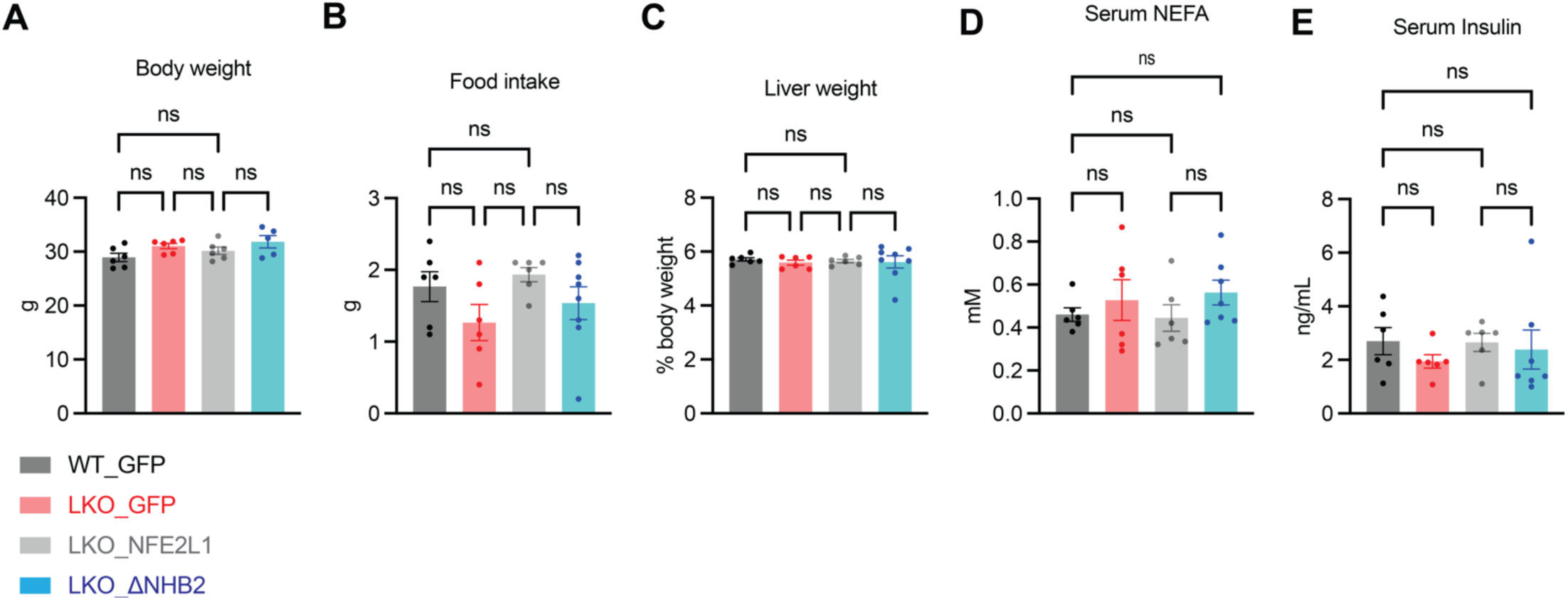
AAV-mediated NFE2L1 expression does not alter metabolic parameters in mice. 8-week-old NFE2L1 wild-type (WT) or liver-specific knockout (LKO) mice were injected with AAV-GFP, AAV-NFE2L1-WT, or AAV-NFE2L1-ΔNHB2. Three weeks post-injection, mice were fasted overnight and refed for 6 hours (n=6/group). **(A)** Body weight. **(B)** Food intake during refeeding. **(C)** Liver weight. **(D)** Serum non-esterified fatty acid (NEFA) levels. **(E)** Serum insulin levels. Data are mean ± SEM. Statistical significance was assessed by one-way ANOVA.

**Figure S5.**
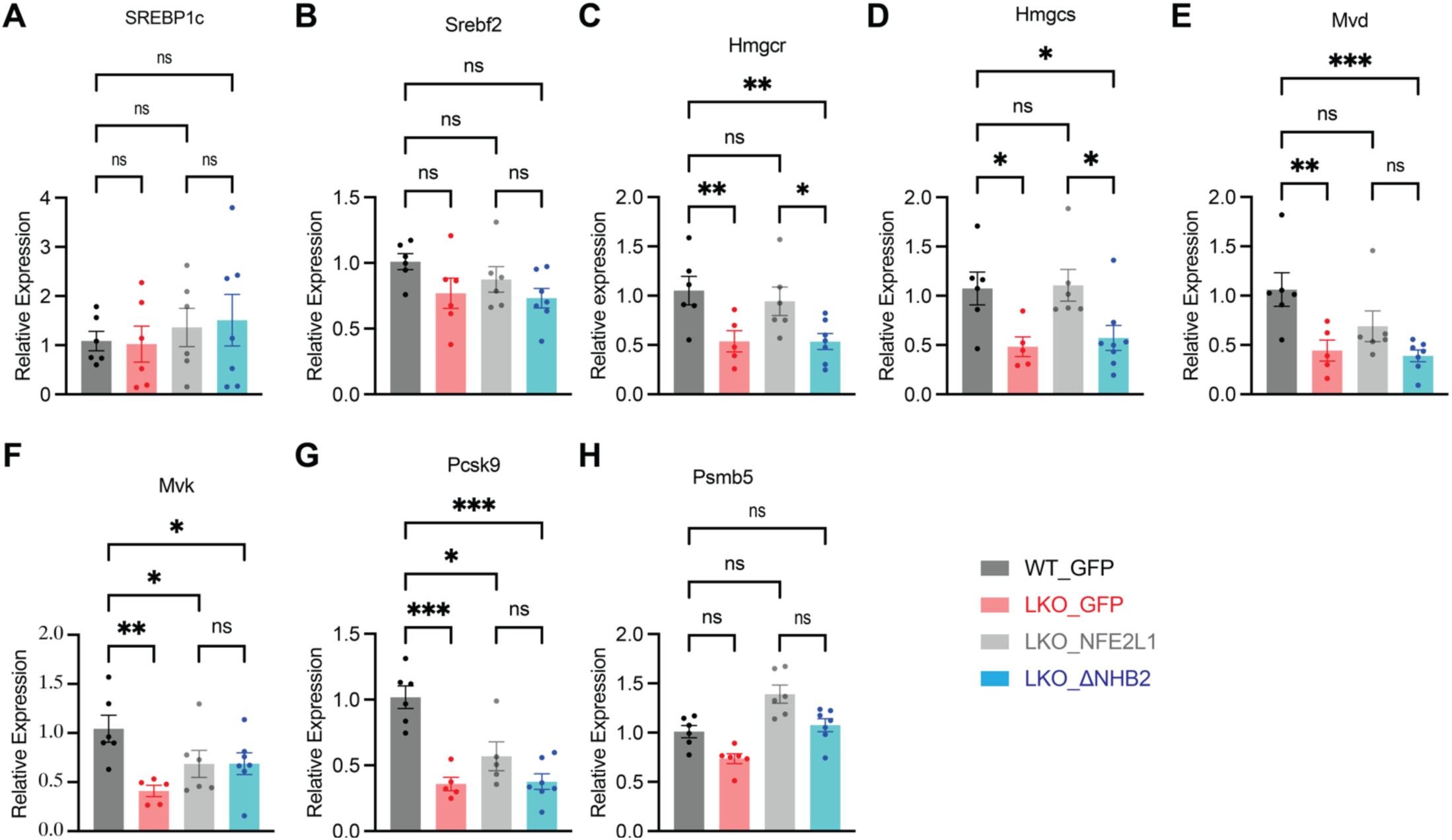
AAV-mediated restoration of NFE2L1 rescues SREBP target gene expression in livers of NFE2L1-LKO mice. 8-week old NFE2L1-WT or -LKO mice were injected with AAV-GFP, AAV-NFE2L1-WT or AAV-NFE2L1-ΔNHB2. Three weeks after injection, mice were fasted overnight then refed for 6 hours (n=6/group). **(A-H)** Relative mRNA levels of SREBP target genes in refed livers. Expression of the genes were normalized to 18S. Data are the means±SEM. Statistical significance was assessed by one-way ANOVA.

**Figure S6.**
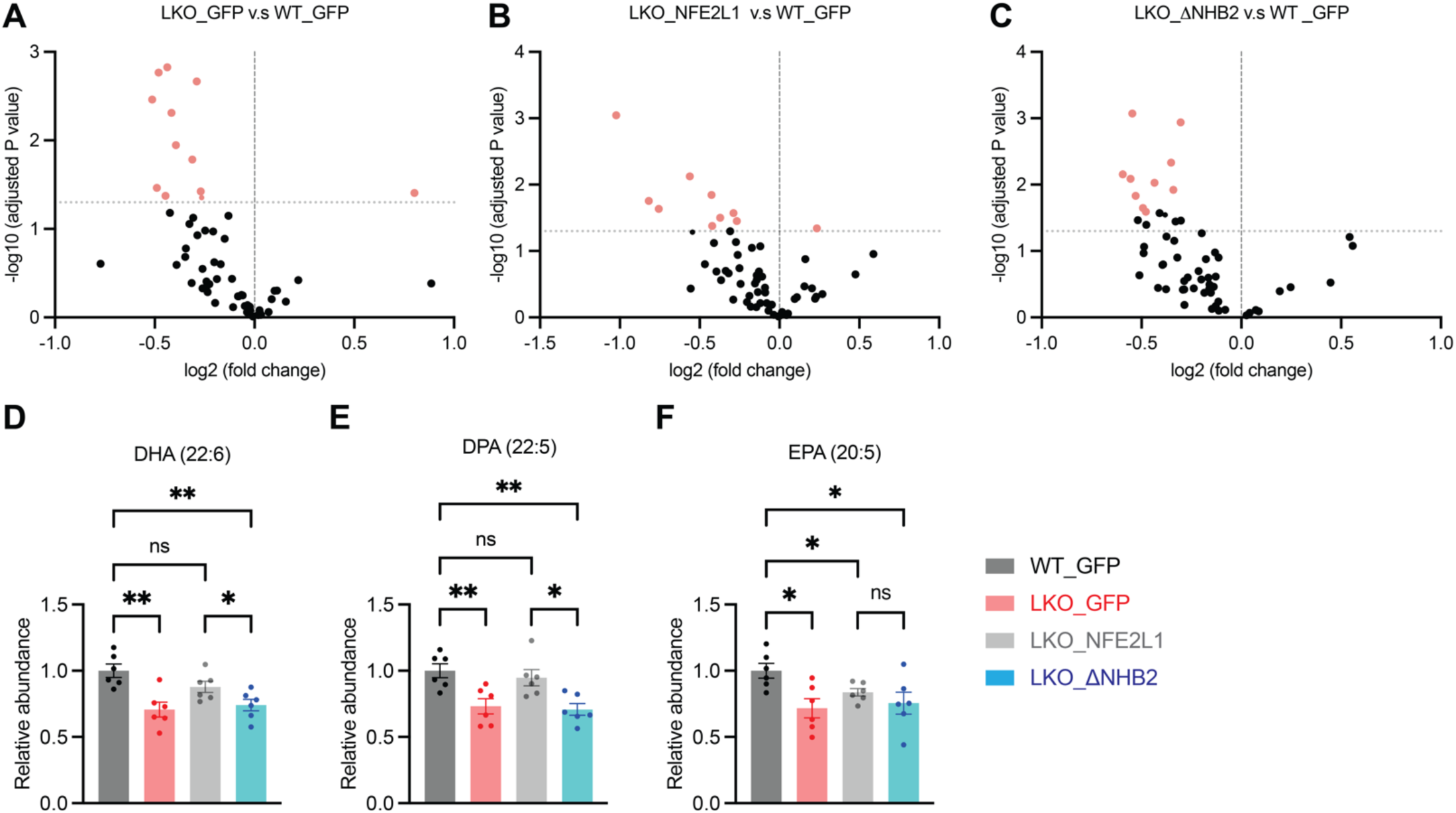
Effect of NFE2L1 -INSIG1 axis on serum phosphatidylcholine (PC) and PUFAs. (A-C) Serum lipidomic analysis for phospholipid species with different fatty acid compositions (n=6 per group). **(D-F)** Serum DHA, DPA and EPA within TG pool. Red indicates p<0.05. Statistical significance was assessed by t-test, *p<0.05.

**Figure S7.**
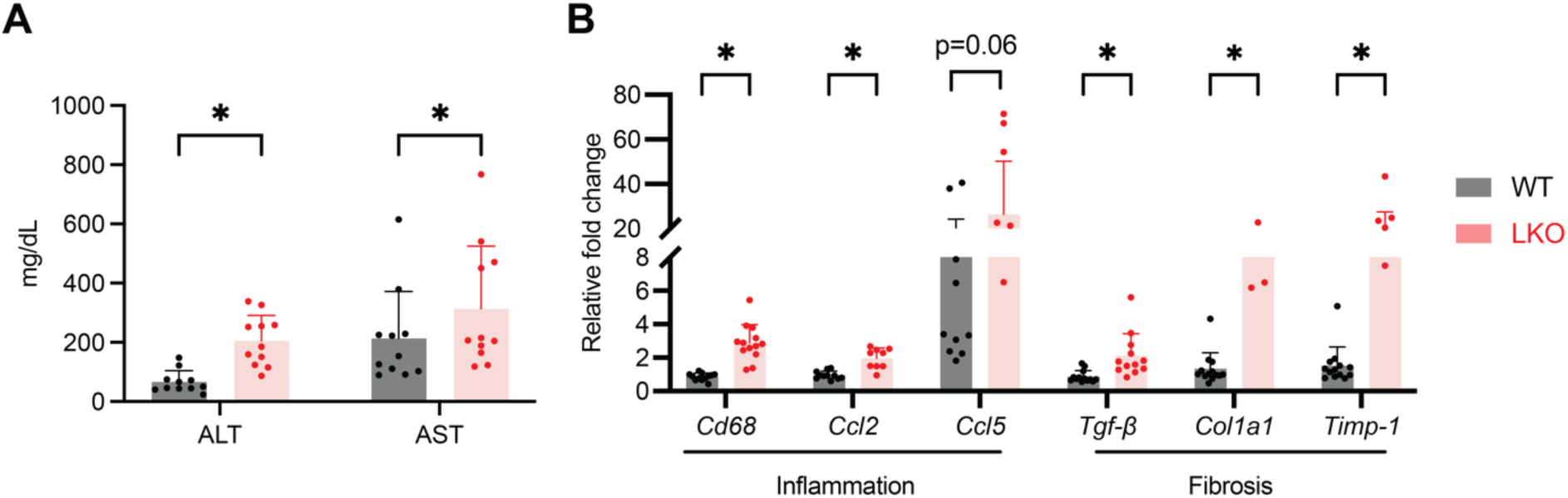
NFE2L1 deficiency exacerbates liver injury, inflammation, and fibrosis. NFE2L1-WT or -LKO mice (8-week old) mice were either fasted for 16 hours or fasted for 16 hours then refed for 6 hours (n=11/group). **(A)** Serum ALT and AST. **(B)** Relative mRNA levels of genes involved in inflammation and fibrogenesis. Total RNA was extracted from livers and subjected to quantitative real-time PCR analysis. Expression of the genes were normalized to 18S. Data are the means±SEM. Statistical significance was assessed by t-test, *p<0.05.

---

**Table S1.** Lipidomic Profiling of Serum Triglycerides Species.

| Compared to WT_GFP mice |  |  |  |  |  |
| --- | --- | --- | --- | --- | --- |
| LKO_GFP |  | LKO_NFE2L1 |  | LKO_NHB2 |  |
| TG species | -Log10 | TG species | -Log10 | TG species | -Log10 |
| TG(20:0_18:1_22:6) | 3.804 | TG(O-18:0_16:0_18:1) | 3.089 | TG(20:0_18:1_22:6) | 4.452 |
| TG(18:1_18:2_22:6) | 3.766 | TG(16:0_16:0_18:0) | 2.480 | TG(19:1_18:2_22:6) | 3.790 |
| TG(16:0_18:2_20:5) | 3.584 | TG(16:0_18:2_18:3) | 2.316 | TG(18:1_20:1_22:6) | 3.759 |
| TG(19:1_18:2_22:6) | 3.536 | TG(O-20:0_16:0_18:1) | 2.110 | TG(14:0_16:0_18:0) | 3.523 |
| TG(18:1_19:1_22:6) | 3.460 | TG(O-24:2_16:0_18:1) | 2.055 | TG(18:1_20:4_24:5) | 3.449 |
| d5-TG(40:9_18:0) | 3.177 | TG(10:0_14:0_16:0) | 2.041 | TG(16:0_16:0_18:0) | 3.275 |
| TG(18:1_20:1_22:6) | 3.013 | TG(14:0_16:0_18:0) | 1.929 | TG(15:0_18:1_22:6) | 3.166 |
| TG(16:1_18:1_22:1) | 2.986 | TG(16:0_18:2_20:5) | 1.912 | TG(17:0_18:1_22:6) | 3.026 |
| TG(14:0_15:0_22:6) | 2.890 | TG(15:0_18:1_22:6) | 1.872 | TG(16:1_18:3_18:3) | 2.897 |
| TG(17:0_18:1_22:6) | 2.817 | TG(16:0_18:2_22:6) | 1.769 | d5-TG(40:9_18:0) | 2.668 |
| TG(19:0_18:1_18:2) | 2.803 | TG(O-24:2_16:0_18:0) | 1.718 | TG(16:0_22:0_22:6) | 2.448 |
| TG(16:0_22:0_22:6) | 2.750 | TG(O-16:0_16:0_18:1) | 1.679 | TG(17:0_18:1_24:6) | 2.431 |
| TG(18:1_19:1_18:2) | 2.739 | TG(28:1_9:0)+OO:(s) | 1.560 | TG(16:0_20:0_22:6) | 2.385 |
| TG(18:1_22:1_18:2) | 2.570 | TG(O-20:1_16:0_18:1) | 1.545 | TG(16:0_24:5_22:6) | 2.382 |
| TG(22:0_18:1_18:2) | 2.564 | TG(15:0_18:2_22:6) | 1.534 | TG(15:0_16:1_22:6) | 2.361 |
| TG(21:0_18:1_18:2) | 2.547 | TG(O-16:0_16:0_18:2) | 1.495 | TG(18:1_20:3_22:6) | 2.349 |
| TG(17:1_18:2_22:6) | 2.527 | TG(18:2_22:6_22:6) | 1.481 | TG(16:0_18:2_22:6) | 2.277 |
| TG(16:0_24:1_22:6) | 2.518 | TG(18:2_22:6_24:6) | 1.344 | TG(14:0_15:0_22:6) | 2.269 |
| TG(18:1_22:1_22:6) | 2.514 | TG(18:2_18:2_22:6) | 1.338 | TG(14:0_18:1_18:3) | 2.219 |
| TG(15:0_16:1_22:6) | 2.448 | TG(16:0_17:0_18:0) | 1.335 | TG(17:1_18:1_22:6) | 2.213 |
| TG(18:1_18:1_18:2) | 2.439 | TG(16:1_18:3_18:3) | 1.319 | TG(14:0_18:1_18:2) | 2.156 |
| TG(15:0_18:1_22:6) | 2.420 | TG(14:0_18:1_18:3) | 1.302 | TG(16:1_18:1_22:1) | 2.110 |
| TG(17:0_18:1_24:6) | 2.397 |  |  | TG(19:0_18:1_18:2) | 2.099 |
| TG(16:0_18:1_23:1) | 2.364 |  |  | TG(16:0_24:1_22:6) | 2.079 |
| TG(17:1_18:1_22:6) | 2.313 |  |  | TG(16:0_18:1_22:5) | 2.071 |
| TG(16:0_21:0_22:6) | 2.312 |  |  | TG(16:0_21:0_22:6) | 2.069 |
| d5-TG(36:4_18:2) | 2.281 |  |  | TG(16:0_18:1_22:6) | 2.050 |
| TG(15:0_18:2_22:6) | 2.229 |  |  | TG(16:0_18:2_20:5) | 2.015 |
| TG(16:0_24:5_22:6) | 2.174 |  |  | TG(15:0_18:2_22:6) | 2.005 |
| TG(16:0_24:0_22:6) | 2.170 |  |  | TG(18:0_18:1_22:6) | 1.972 |
| TG(16:0_20:5_22:6) | 2.166 |  |  | TG(12:0_18:1_18:3) | 1.922 |
| TG(16:0_18:1_22:6) | 2.165 |  |  | TG(18:1_22:6_24:6) | 1.908 |
| TG(16:0_18:2_22:6) | 2.157 |  |  | TG(16:0_18:2_18:3) | 1.898 |
| TG(16:1_22:6_22:6) | 2.151 |  |  | TG(17:1_18:2_22:6) | 1.862 |
| TG(18:1_20:1_20:3) | 2.034 |  |  | TG(16:0_20:5_22:6) | 1.853 |
| TG(18:1_22:6_22:6) | 1.896 |  |  | TG(18:2_22:6_24:6) | 1.825 |
| TG(17:1_18:1_18:2) | 1.868 |  |  | TG(18:1_22:5_22:6) | 1.810 |
| TG(22:1_18:2_22:6) | 1.849 |  |  | TG(18:1_22:6_22:6) | 1.750 |
| TG(18:0_18:1_22:6) | 1.849 |  |  | TG(18:2_22:5_22:6) | 1.735 |
| TG(18:0_18:1_18:2) | 1.809 |  |  | TG(16:0_24:0_22:6) | 1.627 |
| TG(16:0_18:1_24:1) | 1.806 |  |  | TG(17:0_18:1_22:5) | 1.616 |
| TG(18:2_18:2_22:6) | 1.804 |  |  | TG(18:1_24:5_22:6) | 1.601 |
| TG(18:2_18:2_20:5) | 1.797 |  |  | TG(17:0_18:1_20:4) | 1.559 |
| TG(16:1_18:3_18:3) | 1.754 |  |  | TG(18:1_18:2_22:6) | 1.543 |
| TG(14:0_16:0_18:0) | 1.750 |  |  | TG(18:1_20:1_20:3) | 1.541 |
| TG(16:0_18:1_22:5) | 1.742 |  |  | TG(14:0_18:2_18:3) | 1.491 |
| TG(16:0_18:2_18:2) | 1.735 |  |  | TG(14:0_16:0_18:3) | 1.473 |
| TG(O-15:2_3:0_18:2) | 1.732 |  |  | TG(27:1_9:0)+OO:(s) | 1.460 |
| TG(16:0_16:0_18:0) | 1.732 |  |  | TG(21:0_18:1_18:3) | 1.431 |
| TG(22:7_16:0_20:5) | 1.714 |  |  | TG(18:1_22:1_18:2) | 1.413 |
| TG(O-18:0_18:1_18:2) | 1.701 |  |  | TG(18:1_20:4_22:6) | 1.396 |
| TG(18:2_22:5_22:6) | 1.689 |  |  | TG(28:0_20:5) | 1.362 |
| TG(15:0_18:1_18:2) | 1.685 |  |  | d5-TG(36:6_20:2) | 1.362 |
| TG(17:0_18:1_20:4) | 1.671 |  |  | TG(10:0_14:0_18:1) | 1.352 |
| TG(17:0_18:1_18:2) | 1.654 |  |  | TG(15:0_16:0_18:0) | 1.342 |
| TG(16:0_20:0_22:6) | 1.647 |  |  | TG(16:0_18:1_18:2) | 1.335 |
| TG(18:1_20:4_22:6) | 1.611 |  |  | TG(15:0_18:1_18:2) | 1.310 |
| TG(18:1_16:0_18:3) | 1.513 |  |  |  |  |
| TG(14:0_18:1_18:2) | 1.491 |  |  |  |  |
| TG(16:0_18:1_22:1) | 1.468 |  |  |  |  |
| TG(12:0_18:1_18:3) | 1.468 |  |  |  |  |
| TG(12:0_18:1_22:6) | 1.448 |  |  |  |  |
| TG(18:1_20:3_22:6) | 1.436 |  |  |  |  |
| TG(18:1_20:1_18:2) | 1.422 |  |  |  |  |
| TG(17:0_18:1_20:1) | 1.414 |  |  |  |  |
| TG(14:0_18:2_22:6) | 1.413 |  |  |  |  |
| TG(14:0_18:1_18:3) | 1.412 |  |  |  |  |
| TG(18:2_18:3_20:5) | 1.395 |  |  |  |  |
| TG(16:0_18:2_18:3) | 1.363 |  |  |  |  |
| TG(O-15:2_3:0_22:6) | 1.361 |  |  |  |  |
| TG(19:0_18:1_20:4) | 1.361 |  |  |  |  |
| TG(23:0_18:1_18:2) | 1.360 |  |  |  |  |
| TG(16:0_18:1_18:2) | 1.350 |  |  |  |  |
| TG(18:1_20:4_24:5) | 1.330 |  |  |  |  |

Note: The table includes only TG species meeting statistical significance threshold $[(-\log_{10}(\text{adjusted } p \text{ value})) > 1.3, \text{ equivalent to } p < 0.05]$

**Table S2.** Lipidomic Profiling of FA species within Serum Triglycerides Pool.

| Compared to WT_GFP mice |  |  |  |  |  |
| --- | --- | --- | --- | --- | --- |
| LKO_GFP |  | LKO_NFE2L1 |  | LKO_NHB2 |  |
| FA species | -Log10 | FA species | -Log10 | FA species | -Log10 |
| (22:6) | 2.537 | (O-18:0) | 2.930 | (22:5) | 2.744 |
| (23:1) | 2.364 | (O-20:0) | 2.110 | (24:5) | 2.735 |
| (18:2) | 2.226 | (O-24:2) | 2.051 | (20:3) | 2.714 |
| (22:1) | 2.218 | (20:5) | 1.580 | (24:6) | 2.692 |
| (22:5) | 2.196 | (O-20:1) | 1.545 | (22:6) | 2.527 |
| (22:0) | 2.129 | (O-16:0) | 1.531 | (18:3) | 2.012 |
| (20:3) | 2.104 |  |  | (9:0+OO:(s)) | 1.978 |
| (24:5) | 2.007 |  |  | (22:0) | 1.680 |
| (20:5) | 1.971 |  |  | (20:5) | 1.481 |
| (24:1) | 1.743 |  |  | (27:1) | 1.460 |
| (3:0) | 1.726 |  |  | (18:2) | 1.444 |
| (O-15:2) | 1.726 |  |  | (28:0) | 1.362 |
| (22:7) | 1.714 |  |  | (18:1) | 1.295 |
| (19:0) | 1.614 |  |  |  |  |
| (24:6) | 1.565 |  |  |  |  |
| (21:0) | 1.429 |  |  |  |  |
| (18:1) | 1.384 |  |  |  |  |
| (19:1) | 1.360 |  |  |  |  |

Note: The table includes only FA species meeting statistical significance threshold [(-Log10(adjusted p value)>1.3, equivalent to p<0.05]

